# Expression of tardigrade intrinsically disordered proteins in *C. elegans* reveals variable effects on aging and stress

**DOI:** 10.64898/2026.10.26.762026

**Authors:** Anna Entin, Daniel Hicks, Elijah A. Roditi, Juri Kim, Aeowynn J Coakley, Adam J Hruby, Athena Alcala, Naibedya Dutta, Maxim Averbukh, Shivay Parihar, Rebecca Aviles Barahona, Thomas J Lee, Veronica Kuo, Melissa Peruch, Vinaya Sahasrabuddhe, Bharat Vivan Thapa, Jamie D Kraft, Peter J Mullen, Gilberto Garcia, Ryo Higuchi-Sanabria

## Abstract

Tardigrades survive extreme environmental stress by expressing intrinsically disordered proteins that stabilize cellular structures during cryptobiosis. Because these proteins protect cells from desiccation, oxidative stress, and DNA damage, they have attracted interest as potential therapeutics to enhance stress resilience in other organisms. However, most studies have relied on cultured cells or germline expression, leaving their effects in somatic tissues largely unexplored. Here, we generated *Caenorhabditis elegans* strains expressing four tardigrade intrinsically disordered proteins, Damage Suppressor (DSUP), cytoplasmic-abundant heat-soluble (CAHS), mitochondrial-abundant heat-soluble (MAHS), and Ramazzottius varieornatus late embryogenesis abundant (RvLEAM), throughout somatic tissues. Surprisingly, constitutive expression of each protein reduced lifespan, impaired motility and reproduction, and failed to improve resistance to most environmental stressors. DSUP toxicity was tissue- and developmental stage-dependent, indicating that both spatial and temporal expression influence its physiological effects. Despite these detrimental phenotypes, several proteins reduced protein aggregation during desiccation-associated stress, demonstrating context-dependent protective effects. Together, our findings reveal that tardigrade stress proteins impose physiological tradeoffs in vivo and underscore the importance of evaluating candidate stress-protective proteins in physiologically relevant somatic models before therapeutic translation.

**Summary Statement:** Constitutive somatic expression of tardigrade stress proteins impairs aging and stress resilience while providing context-dependent protection during desiccation in *C. elegans*.

## Introduction

Tardigrades are hydrophilous microinvertebrates that possess remarkable stress resilience, capable of surviving heat, cold, desiccation, and oxidative stress by entering a cryptobiotic “tun” state. This state involves dramatic physiological changes, including metabolic arrest (Smythers et al., 2024) and upregulation of protective proteins known as Tardigrade Disordered Proteins (TDPs), which stabilize cellular structures (Boothby et al., 2017; Hesgrove and Boothby, 2020; Lim et al., 2024). These proteins, including cytoplasmic-abundant heat-soluble protein (CAHS), mitochondrial-abundant heat-soluble protein (MAHS), *Ramazzottius varieornatus* mitochondrial late embryogenesis abundant protein (RvLEAM), and the chromatin-binding Damage Suppressor protein (DSUP), stabilize macromolecules and cellular structures during physiological collapse and rehydration, enabling tardigrades to survive stresses lethal to almost all other metazoans.

Although TDPs were first identified as essential proteins to survive cryptobiosis in tardigrades, their protective properties extend across phylogeny. Expression of DSUP, MAHS, and RvLEAM in mammalian cell lines revealed distinct differences in stress-protective capacity. Across chronic and acute solvent exposure and multiple cryopreservation assays, DSUP and MAHS consistently showed improvements in cell viability, enhancing survival, preserving metabolic activity, and increasing post-thaw recovery in both 2D and 3D adipose-like U87 cell cultures. In contrast, RvLEAM showed minimal or no improvement, often performing similarly to or worse than the control group (Pimentel, 2023). The effects of TDPs, however, are context dependent. Neurons expressing DSUP exhibit signs of neurotoxicity, including neurite beading, blebbing, retraction, and significantly elevated cell death. DSUP expression drove chromatin condensation, nuclear disruption, DNA double-strand breaks, and γH2AX accumulation, suggesting that the same chromatin binding properties of DSUP that may be protective in one cell type may instead trigger genotoxic stress in others (Escarcega et al., 2023). For example, work in *S. cerevisiae* showed that DSUP can localize directly to nucleosome-rich regions and modulate beneficial transcriptional responses under oxidative stress, further supporting a dual role for TDPs in both physical DNA shielding and transcriptional reprogramming (Aguilar et al., 2025).

Most recently, while this manuscript was in preparation, one study identified that DSUP can also have positive effects when overexpressed in the germline of *C. elegans*. Germline DSUP expression significantly increased resistance to X-ray-induced DNA damage, reducing embryonic lethality across generations and inducing functional, whole-organism genotoxic resistance. This protection mediated by DSUP was not specific to X-ray-induced DNA damage, as animals also showed increased resistance to multiple oxidative stressors, including paraquat, hydrogen peroxide, and hydroxyl radicals. Furthermore, DSUP expression resulted in extended lifespan in *C. elegans*, suggesting a general increase in health (Richaud et al., 2025).

Investigations in whole animals have provided some of the clearest evidence that TDPs like DSUP can functionally enhance organismal stress tolerance and longevity. Therefore, we asked whether TDPs in general could affect lifespan and healthspan when expressed directly in the somatic tissues of *C. elegans*. To determine the impact of diverse TDPs on aging parameters, we created transgenic animals with overexpression of CAHS, MAHS, DSUP, and RvLEAM to better understand how distinct proteins that localize to various domains of the cell can impact whole animal health, stress resilience, and longevity. To comprehensively evaluate the impact of TDP overexpression on stress resilience, aging, and healthspan, we performed a comprehensive series of physiological, cellular, and behavioral phenotypes. Collectively, these assays provide a robust framework for determining how TDP expression influences the trajectory of aging and stress tolerance in a genetically tractable metazoan.

## Results

Previous work has shown that germline overexpression of the TDP, DSUP, can improve stress resistance and extend lifespan (Richaud et al., 2025). However, the translational relevance of germline expression is inherently limited because therapeutic strategies, especially in aging research, generally target somatic tissues rather than the germline. Moreover, germ cells differ fundamentally from somatic tissues in chromatin organization, genome maintenance, and stress-response pathways (Cao and Pocock, 2022; Shin and Mello, 2003), raising the possibility that TDP function and effects may differ dramatically in somatic tissue. We therefore sought to determine whether somatic expression of TDPs could improve stress resilience and organismal health in *C. elegans*.

### Somatic DSUP expression exhibits toxic effects

We first expressed DSUP specifically in somatic tissues using the whole-worm promoter, *sur-5p* (Gu et al., 1998), and confirmed nuclear expression (**Fig. S1A**). Shockingly, we found that somatic cell expression of DSUP resulted in a significant reduction in lifespan (**Fig. 1A**). Because bacterial infection is a major cause of death in *C. elegans* (Zhao et al., 2017), we next asked whether this phenotype reflected altered susceptibility to bacterial pathogenesis rather than accelerated organismal aging. To address this, we repeated lifespan assays using kanamycin-killed bacteria (De Magalhaes Filho et al., 2018), thereby eliminating bacterial proliferation while maintaining a normal food source. DSUP expression remained significantly detrimental to lifespan under these conditions (**Fig. S1B**), indicating that the reduced longevity is not solely explained by increased susceptibility to bacterial infection.

**Fig. 1.**
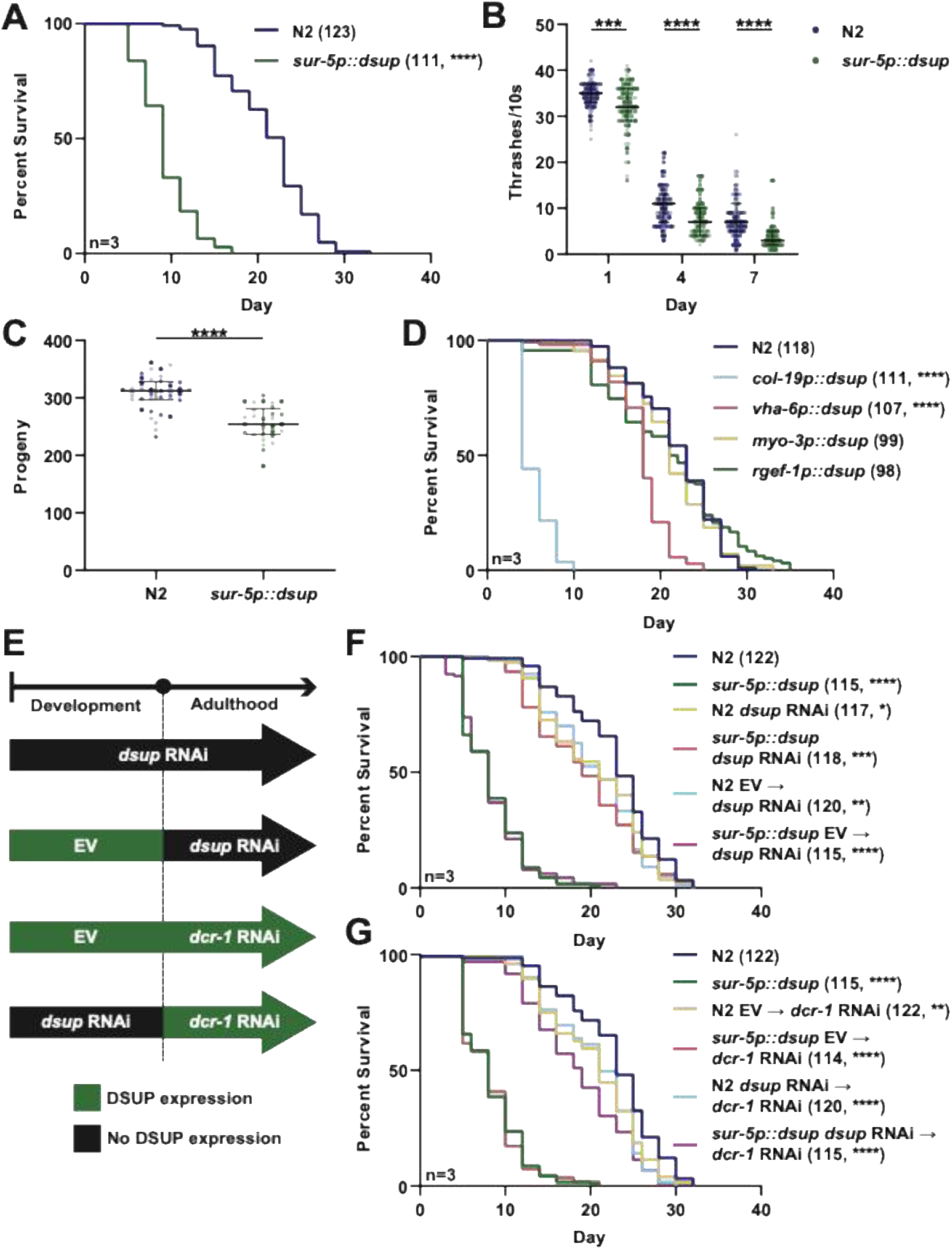
Somatic DSUP expression reduces lifespan and healthspan *C. elegans*. **(A)** Lifespan of wild-type N2 and animals constitutively expressing DSUP in somatic tissues (*sur-5p::dsup*). Animals were maintained at 20°C and survival was scored every two days. Numbers in parentheses indicate total animals scored. Statistical significance was determined by Log-Rank testing. Statistical information for all replicates is available in **Table S2**. Note: the lifespan data was collected and analyzed together with data from Fig. 3A. **(B)** Motility of N2 and *sur-5p::dsup* animals measured by the number of thrashes during a 10-second interval at the indicated days of adulthood. Data is presented as a SuperPlot with each individual dot representing one animal, different shades indicating each independent biological replicate, and lines indicating the median and interquartile range. Statistical significance was determined by non-parametric Mann-Whitney testing. Note: the day 1 thrashing data was collected and analyzed together with data from Fig. 3B. **(C)** Brood size was measured in N2 and *sur-5p::dsup* animals. Data is presented as a SuperPlot with each individual dot representing one animal, different shades indicating each independent biological replicate, and lines indicating the median and interquartile range. Statistical significance was determined by non-parametric Mann-Whitney testing. Note: the brood data was collected and analyzed together with data from Fig. 3C. **(D)** Lifespan of N2 animals and animals expressing DSUP specifically in the hypodermis (*col-19p::dsup*), intestine (*vha-6p::dsup*), muscle (*myo-3p::dsup*), or neurons (*rgef-1p::dsup*). Numbers in parentheses indicate total animals scored. Statistical significance was determined by Log-Rank testing. Statistical information for all replicates is available in **Table S2**. **(E)** Schematic of *dsup* and *dcr-1* RNAi timing. **(F)** Lifespan of N2 and *sur-5p::dsup* animals grown on EV, *dsup* RNAi throughout life, or *dsup* RNAi beginning at day 1 of adulthood. Statistical significance was determined by Log-Rank testing. Statistical information for all replicates is available in **Table S2**. **(G)** Lifespan of N2 and *sur-5p::dsup* animals grown on EV, EV from L1 then switched to *dcr-1* RNAi from day 1 of adulthood, or *dsup* RNAi from L1 then switched to *dcr-1* RNAi from day 1 of adulthood. Numbers in parentheses indicate total animals scored. Statistical significance was determined by Log-Rank testing. Statistical information for all replicates is available in **Table S2**. Note: data from figures 1F and 1G was collected and analyzed together. * = p < 0.05; ** = p < 0.01; *** = p < 0.001; **** = p < 0.0001.

To further assess the impact of DSUP on animal healthspan, we measured locomotor activity using a liquid thrashing assay, which is directly correlated with animal health and remaining lifespan (Herndon et al., 2002). DSUP expression showed a significant reduction in thrashing at all time points (**Fig. 1B**). We also found similar effects on reproductive health, where DSUP expression showed a significant reduction in overall brood size (**Fig. 1C**). To determine whether the effect of DSUP on lifespan had variable effects in distinct tissue, we expressed DSUP specifically in the hypodermis, intestine, muscle, and neurons using cell-type specific promoters. DSUP expression in the intestine or hypodermis resulted in a significant decrease in lifespan, whereas expression in muscle or neurons did not significantly affect lifespan (**Fig. 1D**). These findings suggest that the detrimental effects of DSUP are tissue-dependent, although no beneficial effects were observed in any tissue.

We next asked whether the lifespan reduction was caused by DSUP expression during development, adulthood, or both. To distinguish the effects of developmental and adult DSUP expression, we temporally controlled DSUP knockdown using RNA interference (RNAi). We designed an RNAi construct targeting the transgenic *dsup* sequence and modulated its activity using RNAi against *dcr-1*, which encodes the Dicer enzyme required for processing double-stranded RNA into small interfering RNAs (Knight and Bass, 2001). By sequentially applying *dcr-1* RNAi (Durieux et al., 2011), we inhibited the RNAi machinery and thereby blocked DSUP RNAi, allowing DSUP expression to be restored at defined developmental stages (**Fig. 1E**). To determine developmental effects, we initiated *dsup* RNAi at day 1 of adulthood and found that DSUP expression solely during development was sufficient to significantly reduce lifespan (**Fig. 1F**). In contrast, animals exposed to *dsup* RNAi during development, and moved onto *dcr-1* RNAi at adulthood (i.e., adult only DSUP expression), displayed a mild but significant reduction in lifespan (**Fig. 1G**). Together, these findings indicate that although DSUP expression during adulthood is mildly detrimental, the majority of its negative effect on longevity is established during development.

Previous studies have shown that constitutive expression of TDPs can impair cellular function under basal conditions while enhancing stress resistance or recovery following acute stress, suggesting that these proteins confer context-dependent benefits (Darch et al., 2026; López et al., 2025). Because TDPs are naturally induced during environmental stress in tardigrades, their constitutive expression in unstressed cells may impose physiological costs, which have negative consequences. We therefore asked whether, despite shortening lifespan under basal conditions, DSUP expression could enhance stress resilience in vivo. Consistent with effects on lifespan, we found that DSUP expression resulted in a significant reduction in survival following both heat stress (**Fig. 2A**) and UV-induced DNA damage (**Fig. 2B**). However, we saw no impact of DSUP on survival under oxidative stress (**Fig. 2C**). In contrast, DSUP expression conferred protection during hyperosmotic stress. Hyperosmotic stress was induced by exposure to 500 mM NaCl, a well-established model of desiccation-like stress that rapidly promotes polyglutamine (polyQ) protein aggregation in *C. elegans* (Moronetti Mazzeo et al., 2012). Consistent with this assay, control animals exhibited a robust increase in polyQ aggregates following hyperosmotic stress, whereas DSUP overexpression significantly attenuated stress-induced aggregate formation (**Fig. 2D**).

**Fig. 2.**
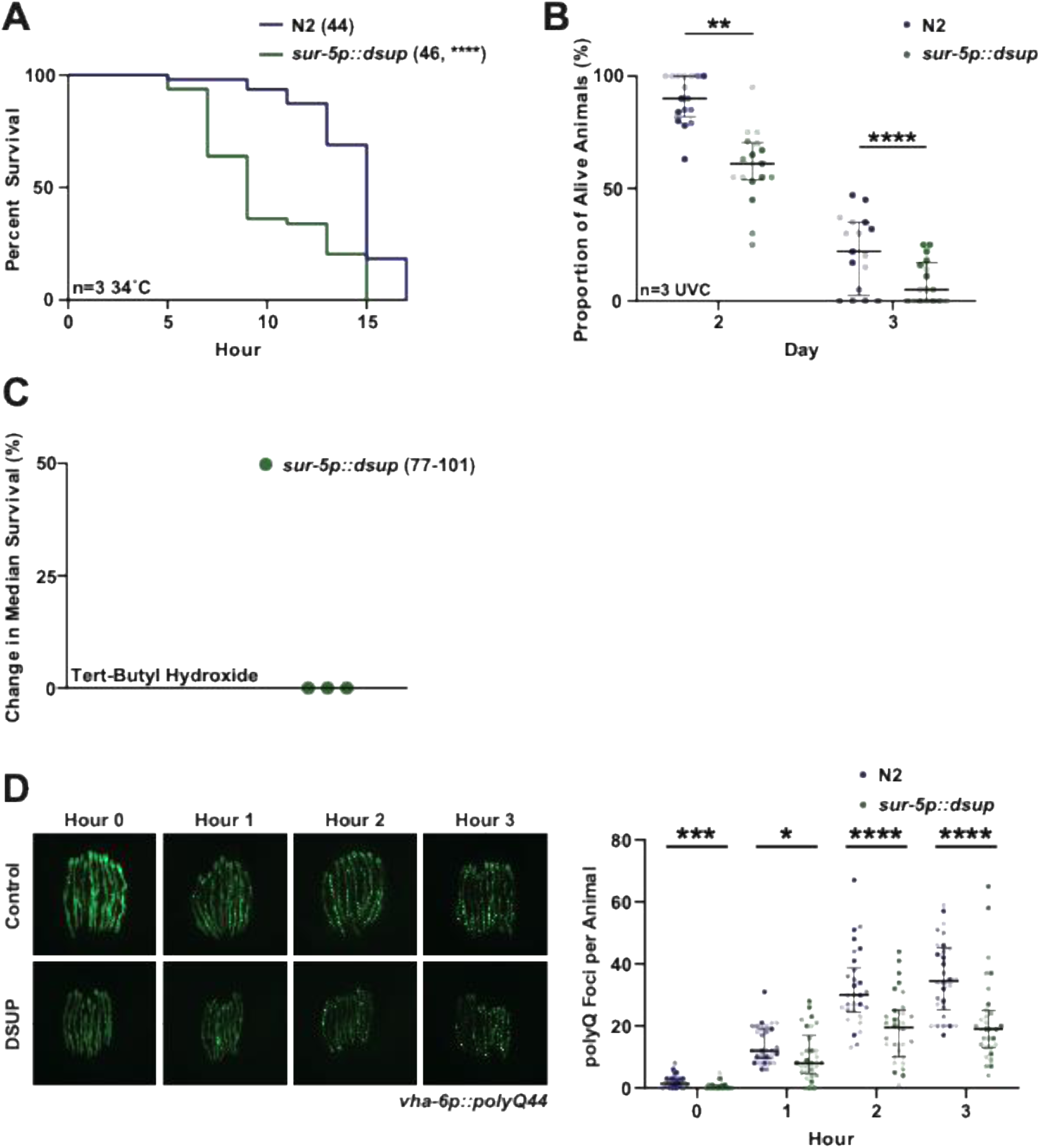
Somatic DSUP expression has stress-specific effects on organismal resilience. **(A)** Thermotolerance of wild-type N2 and animals constitutively expressing DSUP in somatic tissues (*sur-5p::dsup*). Day 1 adult animals were exposed to 34°C and survival was assessed at 2-hour intervals. Numbers in parentheses indicate total animals scored. Statistical significance was determined by Log-Rank testing. Statistical information for all replicates is available in **Table S2**. Note: the thermotolerance data was collected and analyzed together with data from Fig. 4A. **(B)** Survival of N2 and *sur-5p::dsup* animals following UVC-induced DNA damage. Day 1 adult animals were initially exposed to 1 J UVC and subsequently received additional 0.5 J UVC exposures daily. Survival is shown as the proportion of animals alive on the indicated days. Data is presented as a SuperPlot with each individual dot representing one animal, different shades indicating each independent biological replicate, and lines indicating the median and interquartile range. Statistical significance was determined by non-parametric Mann-Whitney testing. Note: the UVC data was collected and analyzed together with data from Fig. 4B. **(C)** Change in median survival of *sur-5p::dsup* animals relative to N2 controls during oxidative stress induced by 6 mM tert-butyl hydroxide. Each point represents one independent biological replicate with N = 77-101. Note: the oxidative stress data was collected and analyzed together with data from Fig. 4C. **(D)** Representative fluorescent images and quantification of polyglutamine (polyQ) foci in control and DSUP-expressing (*sur-5p::dsup*) animals during hyperosmotic stress. Day 1 adult animals expressing *vha-6p::polyQ44* were exposed to 500 mM NaCl and imaged at 0, 1, 2, and 3 hours. PolyQ foci were manually quantified as the number of visible foci per animal. Data is presented as a SuperPlot with each individual dot representing one animal, different shades indicating each independent biological replicate, and lines indicating the median and interquartile range. Statistical significance was determined by non-parametric Mann-Whitney testing. * = p < 0.05; ** = p < 0.01; *** = p < 0.001; **** = p < 0.0001; ns = not significant, p > 0.05.

### Expression of TDPs in C. elegans broadly reduces longevity but has variable effects on stress resilience

Having established that constitutive somatic expression of DSUP impairs lifespan and stress resilience, we next asked whether these phenotypes were unique to DSUP or represented a broader consequence of expressing TDPs in somatic tissues. To address this question, we examined three additional TDPs with distinct predicted subcellular functions: CAHS, MAHS, and RvLEAM. We first confirmed the subcellular localization of each TDP in live animals. Both MAHS and RvLEAM predominantly localized to mitochondria, as demonstrated by colocalization with a mitochondrial matrix-targeted GFP reporter (Kim et al., 2025) (**Fig. S2A-B**). In contrast, CAHS exhibited diffuse fluorescence throughout the cytoplasm without obvious enrichment in any organelle, consistent with its expected cytosolic localization (**Fig. S2C**).

Similar to DSUP, constitutive somatic expression of MAHS, CAHS, and RvLEAM significantly reduced lifespan compared to the wild-type control (**Fig. 3A**). Moreover, the detrimental effects of CAHS and RvLEAM were not based on bacterial pathogenesis, as CAHS and RvLEAM still reduced lifespan on kanamycin-treated bacteria (**Fig. S2D**). However, MAHS expression no longer reduced lifespan on kanamycin-treated bacteria, suggesting that the negative consequences of MAHS may be specific to effects on sensitivity to bacterial infection. Healthspan phenotypes were also broadly impaired across these strains. MAHS, CAHS, and RvLEAM expressing animals exhibited a significant reduction in motility measured by reduced thrashing at all time points (**Fig. 3B**). Moreover, RvLEAM, CAHS, and MAHS expression resulted in a significant reduction in brood size (**Fig. 3C**). Together, these findings indicate that the detrimental effects observed with DSUP are not unique to this protein but are shared across multiple classes of TDPs, suggesting that constitutive somatic TDP expression broadly compromises organismal health.

**Fig. 3.**
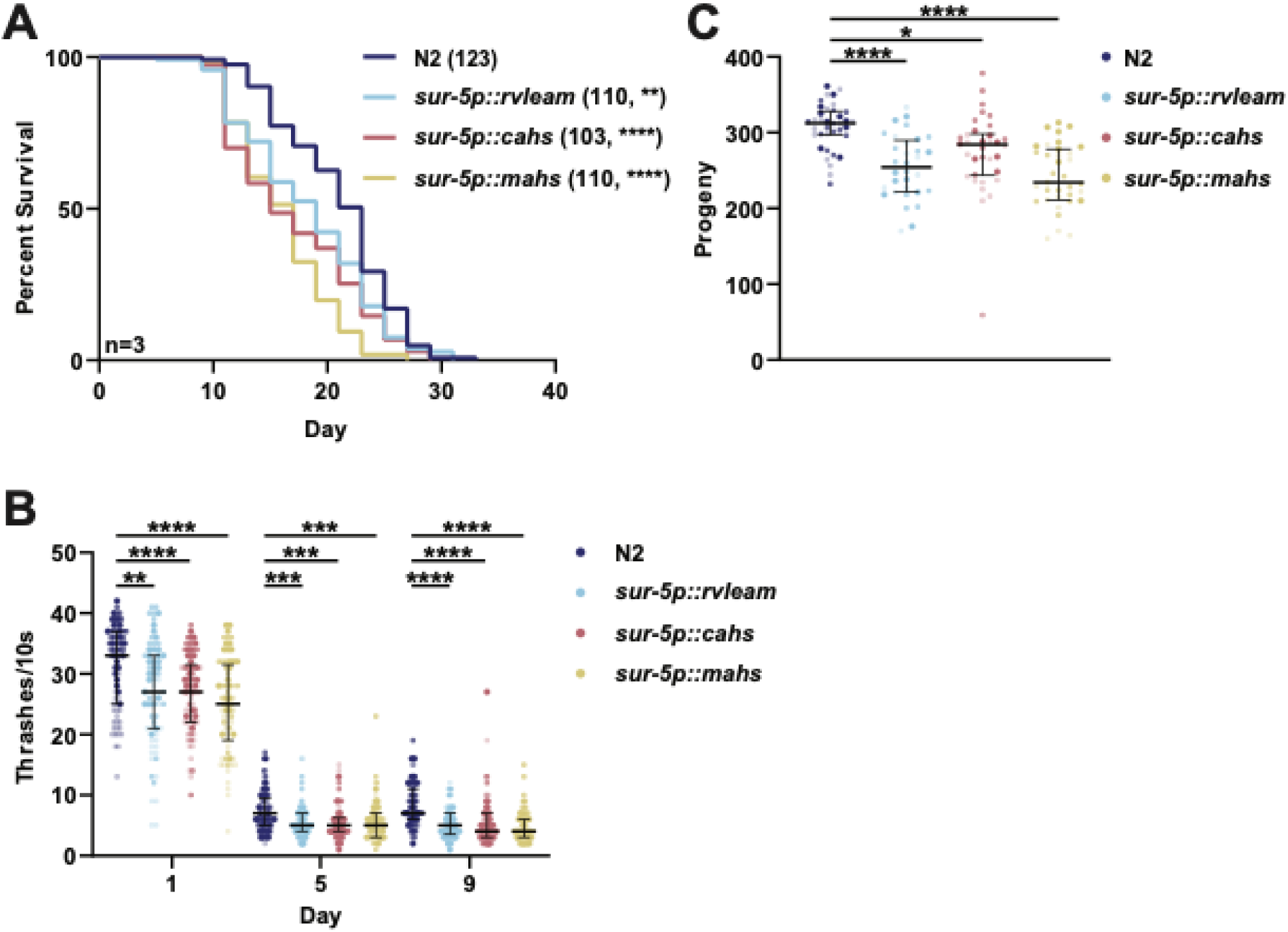
Constitutive somatic expression of RvLEAM, CAHS, or MAHS reduces lifespan and healthspan in *C. elegans*. **(A)** Lifespan of wild-type N2 animals and transgenic animals constitutively expressing RvLEAM (*sur-5p::rvleam*), CAHS (*sur-5p::cahs*), or MAHS (*sur-5p::mahs*) in somatic tissues. Numbers in parentheses indicate total animals scored. Statistical significance was determined by Log-Rank testing. Statistical information for all replicates is available in **Table S2**. Note: the lifespan data was collected and analyzed together with data from Fig. 1A**. (B)** Motility of N2 and RvLEAM, CAHS, and MAHS expressing animals measured by the number of thrashes during a 10-second interval on days 1, 5, and 9 of adulthood. Data is presented as a SupePplot with each individual dot representing one animal, different shades indicating each independent biological replicate, and lines indicating the median and interquartile range. Statistical significance was determined by non-parametric Mann-Whitney testing. Note: the day 1 thrashing data was collected and analyzed together with data from Fig. 1B. **(C)** Total progeny produced by individual N2, RvLEAM, CAHS, and MAHS expressing animals. Data is presented as a SuperPlot with each individual dot representing one animal, different shades indicating each independent biological replicate, and lines indicating the median and interquartile range. Statistical significance was determined by non-parametric Mann-Whitney testing. Note: the brood data was collected and analyzed together with data from Fig. 1C. * = p < 0.05; ** = p < 0.01; *** = p < 0.001; **** = p < 0.0001; ns = not significant, p > 0.05.

One possible explanation for these findings is that constitutive expression of stress-responsive proteins carries an inherent physiological cost. Similar tradeoffs have been reported in other systems, where chronic activation of stress-response pathways enhances survival under stress but compromises growth, reproduction, or organismal fitness under basal conditions (Darch et al., 2026; López et al., 2025). To test this possibility in vivo, we focused on CAHS and MAHS, two heat-soluble TDPs originally identified based on their exceptional solubility following heat treatment and their established roles in tardigrade stress tolerance (Yamaguchi et al., 2012). We therefore expressed CAHS and MAHS under the control of the heat shock-inducible *hsp-16.2* promoter, allowing their expression to coincide with acute heat stress (Link et al., 1999). Surprisingly, inducible expression failed to improve thermotolerance (**Fig. S2E**), suggesting that transient expression of a single TDP is not sufficient to recapitulate the remarkable stress tolerance of tardigrades.

We next asked whether, similar to DSUP, other TDPs modulated stress resilience. In contrast to DSUP, constitutive expression of MAHS, CAHS, and RvLEAM did not significantly alter survival during heat stress (**Fig. 4A**) or following UV-induced DNA damage (**Fig. 4B**). Similar to DSUP, none of these TDPs affected sensitivity to oxidative stress (**Fig. 4C**). However, the effects of TDP expression were context dependent. As observed for DSUP, both CAHS and RvLEAM significantly attenuated desiccation-induced polyQ aggregation, whereas MAHS had no effect (**Fig. 4D**). Together, these findings indicate that although constitutive somatic expression of TDPs fails to broadly enhance organismal stress resistance, select TDPs retain protective activity under desiccation-like stress, consistent with their proposed physiological role in tardigrade cryptobiosis.

**Fig. 4.**
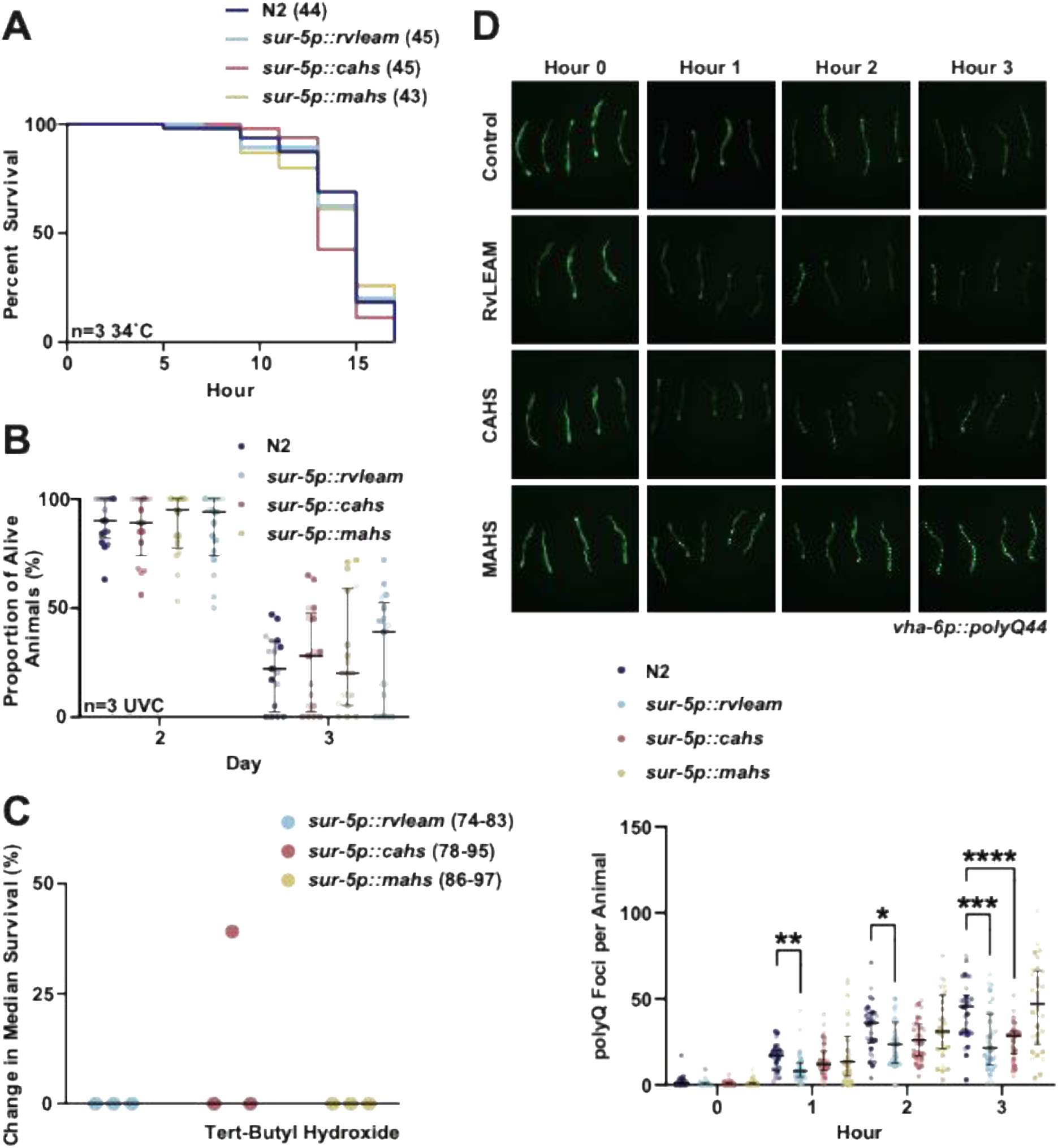
Constitutive somatic expression of RvLEAM, CAHS, MAHS or DSUP has variable effects on stress resilience in *C. elegans*. **(A)** Thermotolerance of wild-type N2 and animals constitutively expressing *sur-5p::rvleam*, *sur-5p::cahs*, or *sur-5p::mahs*. Day 1 adult animals were exposed to 34°C and survival was assessed at 2-hour intervals. Numbers in parentheses indicate total animals scored. Statistical significance was determined by Log-Rank testing. Statistical information for all replicates is available in **Table S2**. Note: the thermotolerance data was collected and analyzed together with data from Fig. 2A. **(B)** Survival of N2, *sur-5p::rvleam*, *sur-5p::cahs*, and *sur-5p::mahs* animals following UVC-induced DNA damage. Day 1 adult animals were initially exposed to 1 J UVC and subsequently received additional 0.5 J UVC exposures daily. Survival is shown as the proportion of animals alive on the indicated days. Data is presented as a SuperPlot with each individual dot representing one animal, different shades indicating each independent biological replicate, and lines indicating the median and interquartile range. Statistical significance was determined by non-parametric Mann-Whitney testing. Note: the UVC data was collected and analyzed together with data from Fig. 2B. **(C)** Change in median survival *sur-5p::rvleam*, *sur-5p::cahs*, and *sur-5p::mahs* animals relative to N2 controls during oxidative stress induced by 6 mM tert-butyl hydroxide. Each point represents one independent biological replicate with N = 74-97. Note: the oxidative stress data was collected and analyzed together with data from Fig. 2C**. (D)** Representative fluorescent images and quantification of polyglutamine (polyQ) foci in control, *sur-5p::rvleam* (RvLEAM), *sur-5p::cahs* (CAHS), and *sur-5p::mahs* (MAHS) animals during hyperosmotic stress. Day 1 adult animals expressing *vha-6p::polyQ44* were exposed to 500 mM NaCl and imaged at 0, 1, 2, and 3 hours. PolyQ foci were manually quantified as the number of visible foci per animal. Data is presented as a SuperPlot with each individual dot representing one animal, different shades indicating each independent biological replicate, and lines indicating the median and interquartile range. Statistical significance was determined by non-parametric Mann-Whitney testing. * = p < 0.05; ** = p < 0.01; *** = p < 0.001; **** = p < 0.0001.

To investigate the mechanisms underlying the physiological effects of somatic TDP expression, we first pursued a targeted analysis of mitochondrial function. A recent study reported that germline DSUP expression induces mitochondrial fragmentation and reduces oxygen consumption, leading the authors to propose that altered mitochondrial physiology may contribute to lifespan extension (Richaud et al., 2025). We therefore asked whether constitutive somatic expression of TDPs similarly altered mitochondrial physiology. In contrast to germline DSUP expression, somatic expression of DSUP, MAHS, CAHS, or RvLEAM did not significantly affect basal oxygen consumption, as measured by Seahorse respirometry (Haroon and Vermulst, 2019) (**Fig. S3A**). To further assess mitochondrial integrity, we examined mitochondrial morphology in the intestine, hypodermis, and body-wall muscle (Kim et al., 2025). Mitochondrial morphology was quantified using mitoMAPR, an automated image-analysis pipeline that measures mitochondrial length, branching points, object number, and mitochondrial footprint (Zhang et al., 2019). Although mitoMAPR enables unbiased quantification of mitochondrial morphology, skeletonization-based measurements can have limitations in resolving complex mitochondrial networks, particularly under conditions of extensive fusion or fragmentation (Lewis and Marshall, 2023; Viana et al., 2015).

Overall, constitutive TDP expression produced relatively limited changes in mitochondrial morphology across the tissues and ages examined. However, several tissue- and age-specific differences were observed. CAHS expression significantly reduced mitochondrial branching points in the hypodermis at day 7 of adulthood, consistent with increased mitochondrial fragmentation, while no other TDP had an impact on hypodermal mitochondria (**Fig. S3B**). Constitutive TDP expression had no impact on mitochondrial morphology in the muscle (**Fig. S4**). Additionally, MAHS expression resulted in a small but statistically significant reduction in mitochondrial length in the intestine at day 1 of adulthood (**Fig. S5A-B**), while no other TDP had an impact in the intestine (**Fig. S5**). These changes were not accompanied by widespread alterations in other mitochondrial morphological parameters or detectable differences in basal oxygen consumption. Together, these findings suggest that although individual TDPs can induce subtle, context-dependent changes in mitochondrial architecture, constitutive somatic TDP expression does not cause the widespread mitochondrial remodeling or respiratory alterations previously associated with germline DSUP expression.

Because our targeted analyses did not identify overt mitochondrial dysfunction underlying the physiological effects of TDP expression, we next employed an unbiased transcriptomic approach. We performed bulk RNA sequencing to determine how constitutive somatic expression of each TDP alters gene expression and to identify molecular pathways that might contribute to the observed reductions in lifespan, healthspan, and stress resilience. Sequencing libraries exhibited consistently high-quality read mapping (**Fig. S6A-B**). Interestingly, TDP-expressing animals were not heavily distinguishable from wild-type controls by general correlation and principal component analysis (**Fig. S6C-D**). However, despite the lack of overt differences overall across datasets, differential expression analysis revealed significant transcriptional changes in all four TDP-expressing strains. Volcano plots demonstrated distinct patterns of both up- and downregulated genes, with the magnitude and extent of differential expression varying substantially among TDPs (**Fig. 5A-D**). RvLEAM exhibited the largest transcriptional response, with 986 significantly differentially expressed genes (DEGs), followed by MAHS (692 DEGs) and DSUP (650 DEGs), whereas CAHS produced a considerably more restricted response (74 DEGs).

**Fig. 5.**
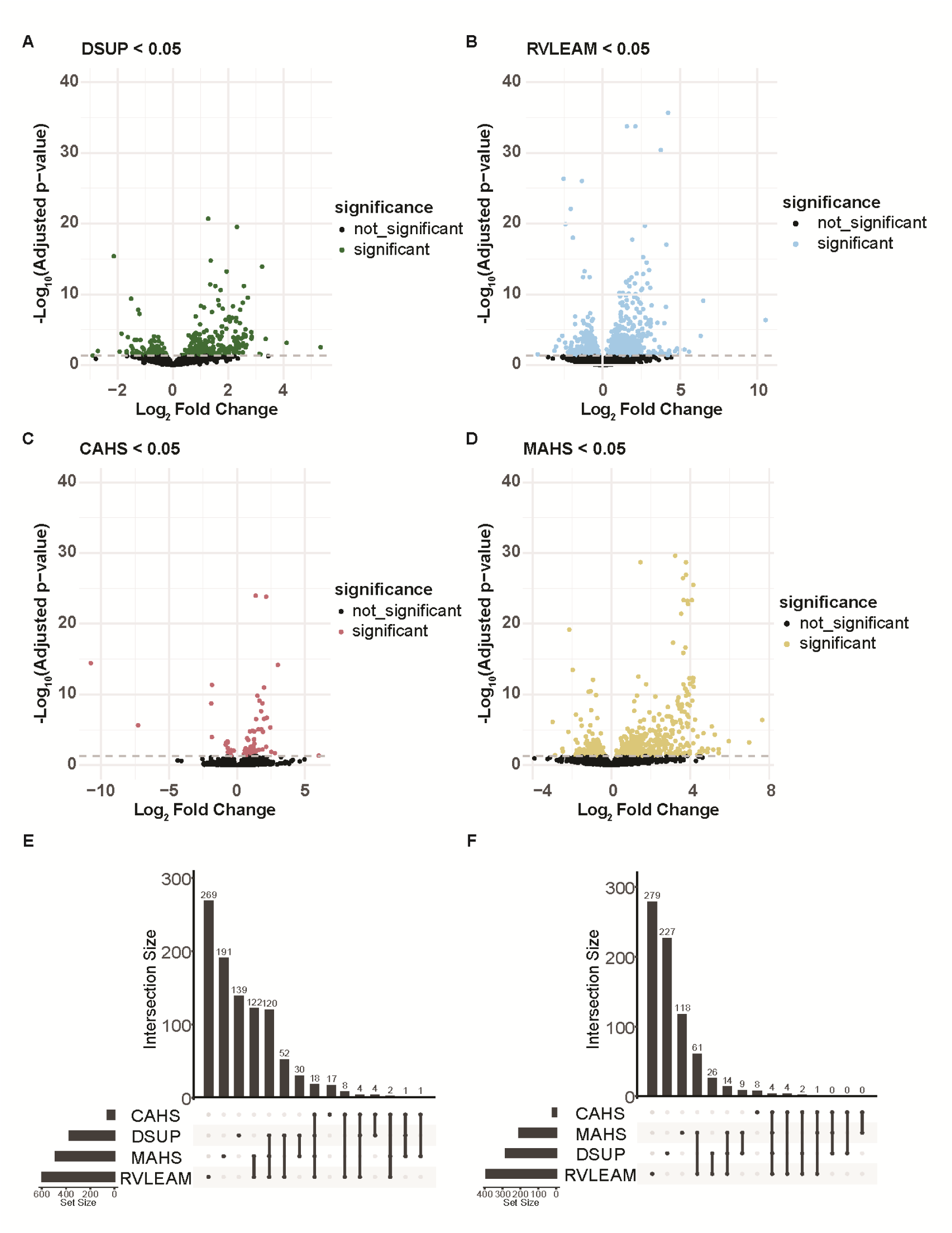
Tardigrade intrinsically disordered proteins induce distinct transcriptional responses in *C. elegans*. **(A-D)** Volcano plots showing differential gene expression in animals constitutively expressing DSUP (A), RvLEAM (B), CAHS (C), or MAHS (D) compared with wild-type N2 animals. Each point represents an individual gene, plotted according to log2 fold change and -log10 adjusted *p*-value. Colored points indicate significantly differentially expressed genes (adjusted *p* < 0.05), and black points indicate genes that did not reach statistical significance. **(E-F)** UpSet plots showing overlap among significantly upregulated (E) and downregulated (F) genes across DSUP-, RvLEAM-, CAHS-, and MAHS-expressing animals. Horizontal bars indicate the total number of differentially expressed genes within each TDP-expressing strain, while vertical bars indicate the number of genes within each intersection. Connected dots below each vertical bar identify the TDP-expressing strains represented in that intersection.

We next asked whether the similar physiological phenotypes produced by the four TDPs were accompanied by a shared transcriptional response. UpSet analysis revealed substantial pairwise overlap among some TDPs, particularly between RvLEAM and MAHS and between RvLEAM and DSUP (**Fig. 5E-F**). However, much of the transcriptional response remained TDP-specific, and only 22 DEGs were shared across all four strains. Thus, despite converging on broadly similar reductions in organismal health and longevity, individual TDPs produced markedly different transcriptional responses rather than a common gene-expression program. To determine whether these TDP-specific transcriptional changes converged at the pathway level, we next performed Gene Ontology (GO) enrichment analysis separately on up-and downregulated genes. DSUP expression strongly induced genes associated with innate immune and defense responses, responses to biotic stimuli, and cuticle development, while downregulated genes were enriched for reproductive and developmental processes, including oogenesis, germ-cell development, gamete generation, and embryonic development (**Fig. 6A, 7A**). Similarly, MAHS expression produced transcriptional signature characterized by induction of cuticle-related pathways, including cuticle development, molting, and extracellular matrix organization (**Fig. 6B**). However, downregulated genes were enriched for chromatin organization, transcriptional regulation, and developmental processes (**Fig. 7B**). RvLEAM expression also induced genes associated with defense responses and cuticle development, but uniquely also enriched pathways associated with ER stress signaling, including the PERK-mediated unfolded protein response, integrated stress response, and ER-nucleus signaling, suggesting RvLEAM alters stress responsive pathways (**Fig. 6C**). No significantly enriched GO terms were identified among RvLEAM-downregulated genes. Finally, CAHS produced comparatively limited pathway-level enrichment, with no significant GO terms among upregulated genes. However, downregulated genes were enriched for innate immune and bacterial-defense pathways (**Fig. 7C**).

**Fig. 6.**
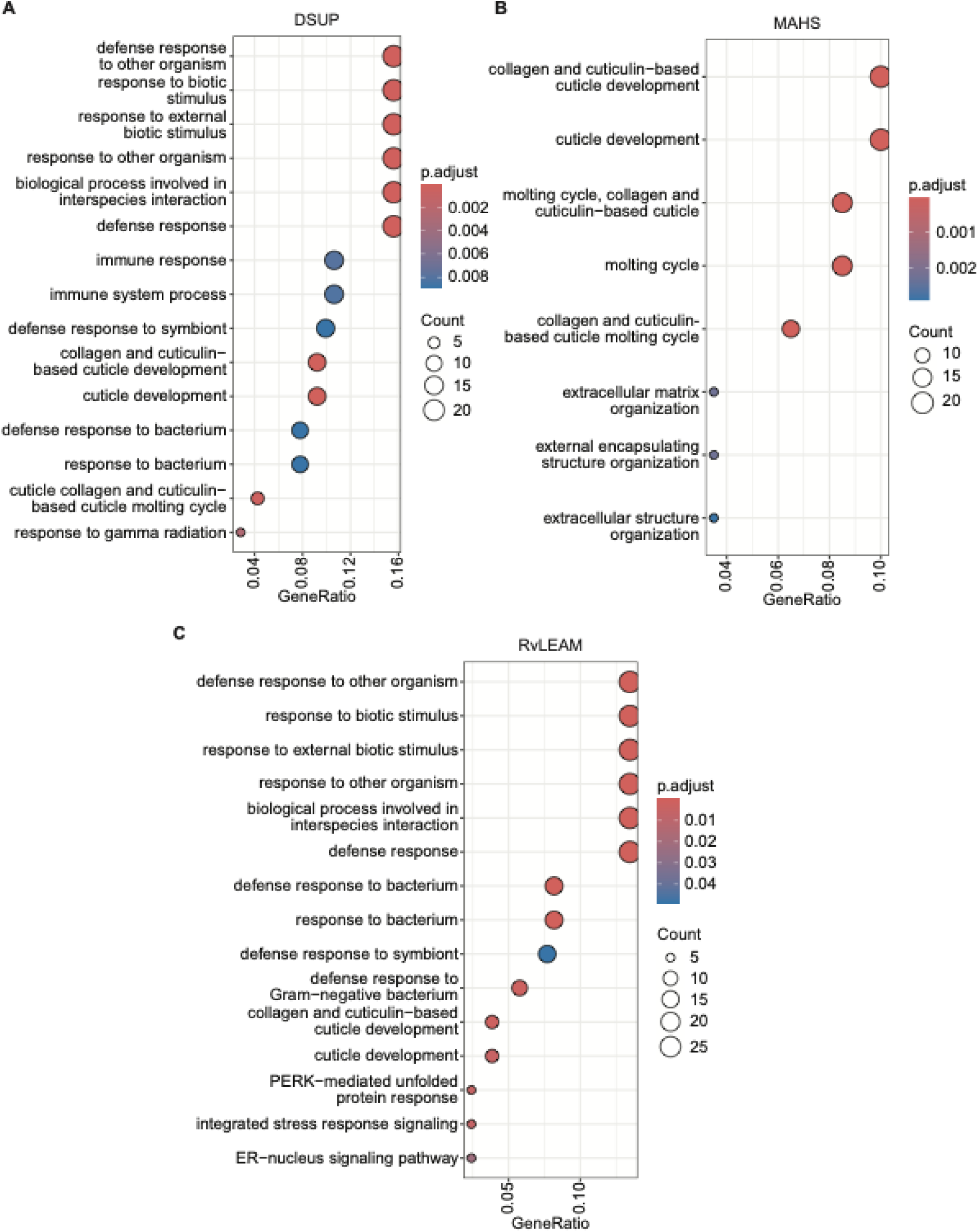
Gene Ontology enrichment analysis of genes upregulated by tardigrade intrinsically disordered protein expression. Gene Ontology (GO) Biological Process enrichment analysis of significantly upregulated genes in animals constitutively expressing DSUP **(A)**, MAHS **(B)**, or RvLEAM **(C)** compared with wild-type N2 animals. Dot size represents the number of differentially expressed genes associated with each GO term, color indicates the adjusted *p*-value, and GeneRatio represents the proportion of genes within the input gene set associated with each term. No significantly enriched GO Biological Process terms were identified among genes upregulated by CAHS expression.

**Fig. 7.**
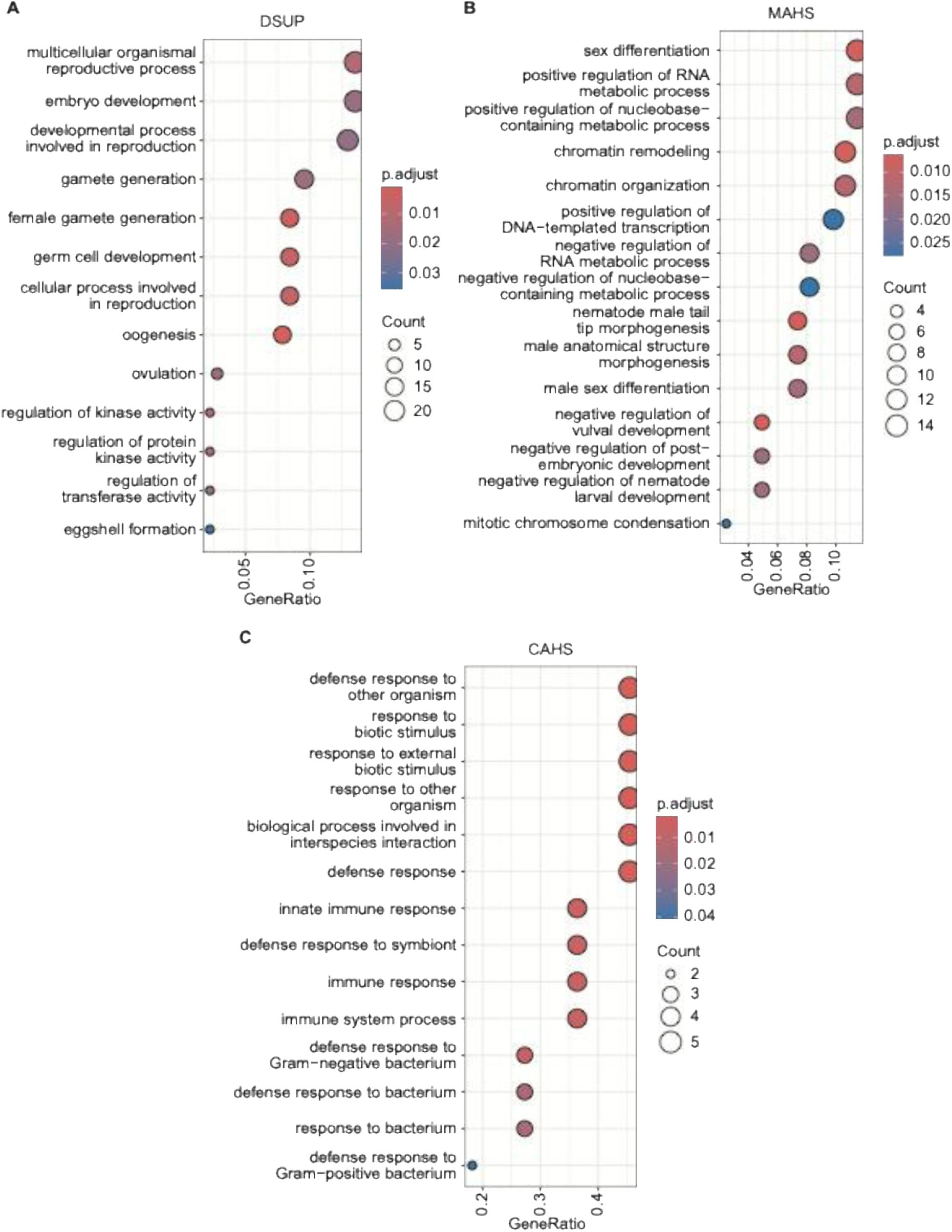
Gene Ontology enrichment analysis of genes downregulated by tardigrade intrinsically disordered protein expression. Gene Ontology (GO) Biological Process enrichment analysis of significantly downregulated genes in animals constitutively expressing DSUP **(A)**, MAHS **(B)**, or CAHS **(C)** compared with wild-type N2 animals. Dot size represents the number of differentially expressed genes associated with each GO term, color indicates the adjusted *P*-value, and GeneRatio represents the proportion of genes within the input gene set associated with each term. No significantly enriched GO Biological Process terms were identified among genes downregulated by RvLEAM expression.

Together, these analyses reveal that TDP expression does not produce a single conserved transcriptional program. Although several biological processes, including immune defense, cuticle remodeling, and developmental pathways, were enriched across multiple TDP-expressing strains, this convergence at the pathway level did not necessarily reflect a common underlying transcriptional response. For example, closer examination of immune-response genes revealed largely distinct patterns of differential expression across individual TDPs, with only a subset of genes similarly affected across multiple strains (**Fig. S7A**). Thus, even when TDPs alter similar biological processes, the specific transcriptional responses underlying these pathway-level signatures remain largely TDP-specific.

Notably, none of the TDPs produced a coordinated transcriptional induction of canonical mitochondrial unfolded protein response (UPR^mt^) or oxidative stress-response genes (**Fig. S7B-C**). Although individual genes within these pathways were altered in some TDP-expressing strains, there was no consistent activation of either program across TDPs. This finding is particularly relevant in light of the mitochondrial fragmentation, reduced oxygen consumption, and enhanced oxidative stress resistance reported following germline DSUP expression, which have been proposed to contribute to its lifespan-extending effects (Richaud et al., 2025). Together with the absence of changes in mitochondrial morphology or basal oxygen consumption in our somatic models, these data indicate that constitutive somatic TDP expression does not recapitulate the mitochondrial or oxidative stress adaptations associated with germline DSUP expression.

## Discussion

TDPs have attracted considerable attention because of their remarkable ability to protect biological structures from extreme environmental stress (Hesgrove and Boothby, 2020). Here, we investigated whether constitutive somatic expression of four representative TDPs, DSUP, MAHS, CAHS, and RvLEAM, could similarly improve organismal health and stress resilience in *C. elegans*. Contrary to expectations, expression of every TDP shortened lifespan and impaired multiple measures of healthspan while providing little protection against most stressors examined in this study. Instead, protective effects were largely restricted to desiccation-like stress, where several TDPs reduced polyQ aggregation. Together, these findings demonstrate that the physiological effects of TDPs are highly context dependent and reveal that constitutive somatic expression produces fundamentally different outcomes than those previously reported following germline expression.

Our findings also emphasize the importance of evaluating TDPs in the appropriate biological context. Previous studies demonstrating beneficial effects of DSUP primarily relied on cultured cells or germline expression (Darch et al., 2026; Hashimoto et al., 2016; López et al., 2025; Richaud et al., 2025). Although these systems clearly showed that TDPs possess intrinsic stress-protective properties, they do not address whether those benefits extend to somatic tissues in vivo, which represent the relevant target for virtually all therapeutic applications. Germ cells differ substantially from somatic tissues in chromatin organization, developmental state, DNA repair pathways, and proteostasis, raising the possibility that TDP function is inherently cell-type dependent. Our results strongly support this idea. Rather than extending lifespan and increasing stress resistance as reported for germline DSUP expression, constitutive somatic expression produced the opposite phenotype, indicating that beneficial effects observed in one tissue cannot necessarily be generalized across the organism.

The tissue-specific effects of DSUP further underscore the context-dependent nature of TDP activity. Previous studies have shown that although DSUP protects transformed mammalian cell lines from genotoxic stress, expression in primary mammalian neurons instead induces chromatin condensation, DNA damage, neurite degeneration, and cell death, demonstrating that DSUP can exert markedly different effects depending on cellular context (Escarcega et al., 2023). In contrast, our studies showed that neuronal expression of DSUP in *C. elegans* did not significantly alter lifespan, whereas expression in the intestine and hypodermis was sufficient to markedly shorten longevity. These observations are not necessarily contradictory. Whereas the mammalian study examined cell-autonomous neuronal toxicity, our study assessed organismal lifespan, which reflects the integrated function of multiple tissues. Together, these findings suggest that DSUP toxicity is highly context dependent and that the tissues most susceptible to its effects may differ across organisms and physiological contexts.

Moreover, temporal manipulation of DSUP expression revealed that the majority of its detrimental effects were established during development. Restricting DSUP expression to development was sufficient to markedly reduce lifespan, whereas expression initiated only during adulthood produced a much milder phenotype. Because DSUP directly associates with chromatin (Chavez et al., 2019), prolonged expression during periods of extensive developmental remodeling may interfere with transcriptional programs or genome organization that are less susceptible following completion of development. These findings suggest that both the tissue and developmental timing of DSUP expression are critical determinants of its physiological consequences.

The stress-resistance phenotypes further highlight the context-dependent nature of TDP function. Despite their proposed roles as broadly protective proteins, constitutive expression of DSUP, MAHS, CAHS, and RvLEAM failed to improve survival under oxidative stress, thermotolerance, or UV-induced DNA damage. In contrast, DSUP, CAHS, and RvLEAM consistently reduced polyQ during hyperosmotic stress, suggesting improved maintenance of proteostasis under desiccation-like conditions. This distinction is particularly noteworthy because desiccation represents the native physiological context in which many TDPs evolved (Hibshman et al., 2020). During cryptobiosis, TDPs are transiently induced in response to severe dehydration rather than constitutively expressed under basal conditions. Our findings therefore suggest that although TDPs retain their intrinsic protective properties, these benefits are largely restricted to the environmental conditions for which they evolved and do not translate into generalized improvements in organismal stress resistance.

One possible explanation for these findings is that constitutive expression of stress-responsive proteins carries an inherent physiological cost. Similar tradeoffs have been reported in other systems, where chronic activation of stress-response pathways enhances survival under stress but compromises growth, reproduction, or organismal fitness under basal conditions (Darch et al., 2026; López et al., 2025). Surprisingly, our data indicate that this explanation is incomplete. Restricting expression of CAHS and MAHS to periods of acute stress did not improve thermotolerance, suggesting that simply limiting expression to stress conditions is insufficient to recapitulate tardigrade-like stress resilience in vivo. Instead, the extraordinary stress tolerance of tardigrades may require the coordinated activity of multiple TDPs together with endogenous stress-response pathways, or additional physiological adaptations associated with cryptobiosis that cannot be reproduced by expressing a single protein. Alternatively, the timing, magnitude, or duration of transgene induction achieved in our system may not faithfully mimic the endogenous regulation of TDPs during cryptobiosis. Future studies combining multiple TDPs or engineering broader stress-response programs may therefore be required to fully reconstruct tardigrade-like stress resilience in vivo.

One surprising finding was the absence of evidence supporting mitochondrial dysfunction as the primary driver of the observed phenotypes downstream of TDP expression. Germline DSUP expression has previously been associated with mitochondrial fragmentation and reduced oxygen consumption, leading to the proposal that altered mitochondrial physiology contributes to lifespan extension (Richaud et al., 2025). In contrast, constitutive somatic expression of DSUP, MAHS, CAHS, or RvLEAM did not significantly alter basal oxygen consumption and produced only limited, tissue- and age-specific changes in mitochondrial morphology. Specifically, CAHS expression reduced mitochondrial branching in the hypodermis of aged animals, whereas MAHS expression modestly reduced mitochondrial length in the intestine of young adults. These changes were not accompanied by widespread mitochondrial remodeling or coordinated transcriptional induction of canonical UPR^mt^ or oxidative stress-response programs. Importantly, the absence of these coordinated stress responses did not reflect a general lack of transcriptional responsiveness to TDP expression. Each TDP produced significant transcriptional remodeling, including enrichment of immune defense, developmental, extracellular matrix, and, in the case of RvLEAM, endoplasmic reticulum stress-response pathways. However, these responses were largely TDP-specific, with limited gene-level convergence despite broadly similar detrimental effects on lifespan and healthspan. Together, these findings suggest that the physiological consequences of constitutive somatic TDP expression are not associated with a common mitochondrial or oxidative stress-response program and differ from the mitochondrial adaptations proposed to accompany germline DSUP-mediated longevity.

Collectively, our findings demonstrate that the remarkable stress-protective properties of tardigrade proteins are not universally transferable through constitutive somatic expression. Rather than functioning as broadly beneficial cytoprotective factors, TDPs exhibit context-dependent activities that depend on the identity of the protein, the tissue in which it is expressed, the developmental stage, and the type of environmental stress encountered. These results establish whole-animal somatic models as an essential platform for evaluating candidate stress-protective proteins and suggest that successful therapeutic application of TDPs will likely require precise spatial and temporal regulation of expression rather than constitutive overexpression. More broadly, our work illustrates that transferring stress-adaptation mechanisms across species involves complex physiological tradeoffs that are not readily predicted from cell culture or germline studies alone.

## Materials and Methods

### C. elegans strains and maintenance

All strains used in this study were derived from the N2 wild-type strain provided by the Caenorhabditis Genetics Center (CGC) and listed in **Table S1**. Worms were typically maintained at 15 °C on standard Nematode Growth Media (NGM) plates that were seeded with OP50 *E. coli* B strain. The NGM plates contained 1 mM CaCl₂, 12.93 μM (5 µg/mL) cholesterol, 25 mM KPO₄ (pH 6.0), 1 mM MgSO₄, 2.5 mM (0.25% w/v) peptone, and 51.3 mM NaCl. For experimental assays, animals were grown at 20 °C on RNAi plates. These plates consisted of NGM supplemented with 1 µM IPTG and 100 µg/mL carbenicillin and were seeded with HT115 E. coli K strain. The bacteria carried either the pL4440 empty vector (EV) or a pL4440 plasmid encoding double-stranded RNA targeting the gene of interest. To obtain age-synchronized populations, animals were synchronized using a standard bleaching protocol (Castro Torres et al., 2022). Briefly, worms were collected in M9 buffer (22 mM KH₂PO₄, 42.3 mM Na₂HPO₄, 85.6 mM NaCl, 1 mM MgSO₄) and exposed to a solution containing 1.8% sodium hypochlorite and 0.375 M KOH in M9 until only embryos remained. The embryos were then washed four times with M9 and incubated overnight (up to 16 h) in M9 at 20 °C on a rotator, allowing them to hatch and arrest at the L1 stage. The synchronized L1 larvae were subsequently transferred onto RNAi plates.

TDP overexpression strains were synthesized by microinjection of standard *C. elegans* vectors containing a codon-optimized coding sequence for Ramazzottius varieornatus CAHS, DSUP, MAHS, and RvLEAM under cell-type specific promoters. Briefly, CAHS, DSUP, MAHS, and RvLEAM were subcloned into pCFJ356 under the control of *sur-5p* (whole-animal somatic expression), *col-19p* (hypodermis), *myo-3p* (muscle), *rgef-1p* (neuron), or *vha-6p* (intestine). Plasmids were injected at 25ng/µL concentration into *unc-119(ed3)* mutants, and wild-type animals were selected to maintain for array-positive animals. *dsup* RNAi was synthesized by subcloning a fragment of the *C. elegans* codon-optimized DSUP construct used for transgenic construction into the pL440 RNAi vector downstream of the T7 promoter. The *dcr-1* RNAi was isolated from the Ahringer library (Kamath and Ahringer, 2003).

### CAHS

ATGAGTTCTAGACAAAATCAACAGTCTTCTTCACAGCATTCAAGTTCTTCACAACAGGGTGG ACAGGGTGGACAGGGAGTACAGGGTTCGTCATCCTATTCTAGAACTGAAGTACATACGTCC TCAGGTGGTCCAACTATTGGTGGTGCACAACGTACAGTTCCAGTGCCACCGGGCAGTCAT TCAGAAGTTCACGAAGAACGGGAAGTGATTAAACACGGAACGAAGACAGAGTCTGAAACA CATGTTGTAACAGTTCCTGTCACCACATTTGGATCAACTAATATGGAGAGTGTGAGAACTG GATTTACTGTTACACAGGATAAGAATTTAACTGTCGCAGCCCCAAATATTGCAGCCCCTATT CATTCTAATCTTGATCTCAATTTAGGAGGAGGTGCCCGTGCCGAGATCACTGCTGGAACAA CTGTGGATCTGTCAAAGATTCAAAGAAAAGACCTTGGTCCGGAAGAATACGCCCGGTATAA AGCAAAAGTTGAACAGCTCGCGAGACAGGACGAACAGGATGCTGGAATGCGTGCAGCGC AATATCGCGAAGAGGTTGAAAGGGACGCTGAGTTGATACGTCAAATTCTTGAACGTCAGCA TATTCGAGACTTAGAGTTTCGTAAAGAGATGGTCGAAAATCAGGTGAATAGACAGGAACGT GAGATTCAACTTGAAGCGGAATATGCTATGCGTGCATTAGAACTAGAACGTAACGCTGCCA AGGAAGCCCTTGAGTCGGCAAAAGCACAAACAAATGTAAACGTAAAAGTAGAAAGTGCAAT AGGCACGACAGTGTCAAAAGGAGCTATACAAACCTCAGCGGATAAAAGTTCGACTACTAAA ACGGGTCCAACTACGGTGACACAAATAAAGCACACGGAGCAGCATACAGAGCGTCGT GACCCGCCAGTA

### DSUP

ATGGCTTCAACCCATCAGTCTTCTACGGAACCATCATCGACGGGCAAGTCAGAAGAGACAA AGAAGGACGCAAGCCAGGGATCTGGACAGGATAGTAAGAATGTGACAGTCACAAAGGGAA CAGGCTCTTCTGCAACTTCTGCGGCGATCGTTAAAACTGGTGGGAGTCAGGGAAAGGACA GTAGCACAACGGCTGGATCGTCGTCTACGCAAGGTCAAAAGTTTTCGACAACTCCGACTGA TCCTAAGACATTTTCAAGTGATCAGAAAGAAAAGAGTAAGTCACCGGCAAAAGAGGTTCCA TCGGGCGGAGACAGCAAATCTCAAGGGGATACTAAAAGCCAGTCAGATGCGAAGTCATCC GGCCAGTCCCAAGGTCAATCAAAGGATTCTGGAAAGAGCTCATCTGATTCATCAAAATCAC ATAGTGTGATAGGTGCCGTGAAGGATGTTGTGGCTGGGGCTAAGGACGTAGCCGGGAAG GCTGTAGAAGACGCCCCGTCTATTATGCACACCGCCGTTGACGCCGTTAAGAATGCTGCT ACTACAGTTAAAGACGTTGCCTCAAGCGCCGCTTCCACAGTAGCCGAAAAGGTGGTGGAC GCCTATCATAGCGTCGTAGGTGATAAAACTGATGATAAGAAGGAAGGAGAACATTCAGGAG ATAAGAAAGATGATTCTAAGGCAGGTTCAGGATCCGGTCAGGGTGGAGATAATAAGAAATC CGAGGGTGAAACGTCAGGACAGGCTGAGAGTTCATCGGGAAATGAGGGTGCAGCACCTG CTAAGGGACGAGGCAGAGGGAGACCACCTGCTGCGGCCAAGGGCGTGGCAAAAGGAGC AGCCAAAGGAGCGGCTGCATCAAAGGGTGCAAAATCAGGCGCAGAGTCGTCTAAAGGCG GCGAGCAATCCTCGGGTGACATTGAAATGGCTGACGCAAGTTCTAAAGGTGGATCAGATC AAAGAGACTCAGCTGCTACTGTGGGTGAGGGTGGAGCTTCGGGATCCGAAGGCGGTGCG AAGAAGGGAAGAGGAAGGGGAGCCGGGAAGAAGGCAGATGCAGGAGATACTTCGGCCGA ACCGCCCCGAAGAAGTTCAAGACTTACCAGCAGCGGAACTGGAGCCGGGTCTGCCCCGG CCGCTGCAAAGGGTGGCGCCAAAAGAGCAGCAAGCAGCTCAAGCACCCCATCAAATGCCA AGAAACAGGCTACAGGTGGAGCCGGAAAGGCCGCCGCAACAAAGGCCACGGCAGCTAAG TCAGCCGCTAGTAAGGCACCTCAAAACGGAGCTGGAGCTAAGAAGAAAGGTGGTAAAGCA GGCGGCCGAAAACGCAAAGACCCGCCAGTA

### MAHS

ATGAGCCGGTATCTTTTGCGGGATGTTCAAGCAGTCTTAAGAGGTGTAAGAAAGGTCGCTG AATCTAGTCTCAAACTAGAAACCGAAAAGGTATCTCTTCGATTAGGAGATTTCAGATCTCAA CCATCGCTGAGGTCGGTCCCAGCCTCACTTACCTCTCGCTCCCAAGCTTTCAGTCTCCAAG AAATTGCAGCCCGAGCAGGCGTAGTGCTTCGTGGCGTTCAGCAGCAATTTAGAAATGTGA CAGGTGTTAACGCTGCCCCGGTGGTTGCTTTCGACAACGGCTCGGTGCTTTATTCAGAGC GAATTCATAGCCAATCTAGCCAGAAACAAGCACCAACAACTGTCCCTACCGGTAGCGTATC CAACTCTCCCCAGCCAGAGGGGAAAGCTAATGAGGCAGCCGAGAGGGCCAAGCAATTCAT GAACCCACCGGTGGCTCCGATGGACCCGGTTGATAAGAACGAGTTCGTTGCAATGCCAGA AATGGGACGATCTAACGGTAACGGTGAGAATAAGCAGGCGGCCGACTTTATGAAGAATCA GGGAGATACAGACATGGACAGTCAATATGCTCCGGACAGCTCTAAGAATACCAAGTCAGTA CCAACAAAAGAGATTGTGGCAGAGGACGGAAGTATGTCGATAGAAGACATAAAGAAGGCC ACCCAAGTAACACCCGGTGTCGCTGTCAAGAATGAAGGAGTGGACCCGCCAGTA

### RvLEAM

ATGTTCCTTGCTAGAAATGCCGGGCGAGCAGGCTACCGCGGAGTTGTTGCATATCAACAG GCGGCAAGCTTTTCCGTATCATCCGCCAAAGCGGCCGGTTCGAGAAGTTCAGGCGGATCA GATGCAGGAGACTACGCGCGAGAAGCGGCAGAGCACGCAAAGGCCGGCCTTAAAGACCT AAAGAACGAGGCTTCCTGGAAGGCAAAGGGCGTAGCAAATCAGGCTGCCGGTGCTTTTGA GCGGGCCAAAGACACTGTTAAAGAGGGAGTGCACGATATGAAACGATCCGGCTCGCGAGT ATTTGAGCAGGGGCAAGAAGAGGTTGAAGCAGGAGCGCAGCATGCTAAAGCCGGATACCA ATCCGCGAAGAACGCAGCACAGGATACCGCAGCTACACTCAAGGATAAGGCTGGATCCGC GTGGAACCAGGCAAAGCATGTCGTAGAGGACAAAGGCGAAGACGTGGTAGAGGCGGTAA AAGATACCGCCTCAAAGGTGTGGGGAAAGGCTAAGCACGTTGCAGAAGATGTAAAAGAAA ATGCTCAGAGCCCCGGTGGAATAGCTGACAAAGCTTCAGACGTTTGGAGCGCCGCTAAAG ACAAGGCCGCCGACGTTCTCTCTGGTGCAAAACATACCGCAGAAAATCTTGCACATAAGGC TCAGGCAGCAATTCATGACGCCACTGCCAGTAGCGGTTCCCAATCTCAATCACAAAGTCAA AGCCAATACCGGCAGGGTCAGCAGCAAGGCCGACAGGACCAACAACAGTCTAAATCGCAA TGGGGTCAAACCTCTCCACAATCACCTGACGGCTTTAGGCCCCAGGCAGGACAAGGGCCT CAGGGCGGGAAAGGTCCAGGACAGGCCGGTGGACGTAGAGACCCGCCAGTA

### sur-5p

tacatcataacagagaaaactctttaaaatcatattgttcagccaggtcacagtcaaggacaattttgaatttgactcaatttcagttacag gctcggataacggcgctgccacgaccaccctggccccttctgacccaacttgttaaatattaatctaaatacaaataagatttagtatga acatttactgttcattttgaatgttgtttttttacacattttcgtagttgaaaattttgacatattttaagtgttggaaaagatatgagacatttgagc gaactcgtccaaatttgtgaaaatttccaaaaatcatcgtgaaaaattaaaaaacatattttcagagaaaaggcttaaacatttttgaata tgaataagtaaccttttttaaaaatcggcatgtttttttctttgaagatcaatatttttgcgaaagcctacgatactaaaaaatattaacagca aaaaggtgtaaaattaaaaagtgattgtttcatctgttgacaaaagttttgtcaaacaactggttttggaatttatatgcaaaaacctgtaa aaagagtgtttattgaatcttggacttgacttgtttcatcggactgtaatttagcaggggcttaagataggatgaattggtaaagtgataaa atatagctattggtatgcaaaatattgttacaattgattactttttggtatcattcacgtttctgaagttaccgccactcaaagttaaataaaat aatttttgcaagcatttttgacaacttaaacgaatatggcgggaccaaaaaggccatgcctggagctgttattgcatgtttttgataaactta ctgtcatacatcagggtacataaggtttttgatttcacacaaagaaaaaaaaatcgaaaaaaaaacaatttttttccgttgtcatagttga aaaaccctgttgctaaacaattcttctaatttttcgtggtaactgctacacaatcggattaaggaaacctaattttttctcacatatgttggag agcctcgatattcttgaactttttgcattcattgtttaatgtttcttcacatagcttcgtatatattgccagcaatgcaaatcttgagccatcggttg aaatgattttgaaaaggtgctcggcatatctgggctgaaaaaaaacttttgtttcaacgcattaaacacttttcggggttctgttccaatttttt aaatagttgtaacacccgctagaatttccgtttcaccaaagaaatctttgttcacagattattatatatttcttttttatgtattttgtccaggtgat cctaaatgtttaaaaaacgttccagcagcttttagtctatacgaagtatattttttcttattttggtttatcggaggtatgcagttccaaatgttgg attttctaccctactagaaatttcctttagttatacaattttaattcgaagttgtatcttgaaaaggaaaccatcatcaacaacaaataagctt gagactagaattgctaatcgtctgcaccaaatatgaaattcgttgaataaattatataaaacagttgacaatcataaagtattttacatac atatgtatagttaatgtcatattagcatatttagtattgagttaacacgcagaaataccaaagaaaccctgtttaattcatttccatgttttattc gctttaactagggaatgaccagaataagtggagcgatattcaaaaaaaaaaaattgtatcggaaagctgacattctctactataagaa tatgactgaaatttttgcccattcgggctggaaatctgaatggaaattgtttatatgaggggatccatggttgtagttacaattttgagaaaa aagtttaaattttaggggaaaagatgttgaatgaatcattatggtgttggtcgacacattgatcatgctgatatcgatcagaaaaattaaat ttgggcttttatatagtttcgaaaaaacacgtttaaatttttgaaaaattttacgttagggtggaattgaaccccagagtcccaccatgcttgt catccactctatccactcggccatctcgctgttgcaggcagtactaattggggatagtgaaaagagttaacgtagaatttcagacgtaaa aatttcagatttccagcccgaatgggccaaaatttcagtcatattcttatagtagagaatgtcagctttccgatacaatattttttttttgaatat cgctccacttattctggtcattccctagtaagttttttcagttttaccagaaataaaaccactcataaaaactataaactacacatttaaactg taaatgttggaaatcattttgaccatcaaacaatagctatcggaaaatgcaaaggccattccgtgatggcacgctcttcttctatccatcg ctccttttcaacgaaacacacacacaaacgcagactattggaaaagaaaaatatagaaaaaaccttttttctccttgtcccttgtgttgat ctcacttttcagtggacgtgtgcagtgtggaggccagagactgtgctgaccgacggacgcgcaggcggtaaacatacgttgcagtgg agagtgtagtccctcgtcggctctctgtgtatttctttcaatgcctgtgtctcggtctatctttgtctctatctgttttaaaagttttttgcaaagtttca atttcttttttattttctcttctttttttgcgcatttattttttttaaagagcgctgttcttcaattggaaaacagtataacgatcgaaataaaaaaaaa acttgaaatagtatcggaattgagatgggggcaattgaaattgaagacgcaacaaaaacatgtttttatcgtctgaaattatctaaacttt gaatctaacgtgtcaatggctacttttacagtagactttttagagtaatgttaaaaaggataatataggtttttcttcaaactattttgtcacgc cgcatatattcttttcatattccattcgcaaagtgcatgatgtaacaaaaatgaaacgtcataattttttaatctttcacactttttttgccggccg ttaaatcaaaatcaaatcaacttaatccaacacgtgcgggcttgcaattttattacaccggtgtcctacagcgccatgttatcttctttgatta gagatctctcacgcaacatttgacattgaaaattttgtctccgaattgttgtgtacctactttgtcccgatgtcccaaattttcactgataacaa gagatatctcaactgttcttatcacgttgatacgtagaacgttttacactaaaccaaattgcagtccccttctacgaaccgtttgctaattgttt tgcatagaacagtgatcaatgtgttgtgctcactagtcaatgatgttcgtctctcatttttcctacaacttcactgaaaaggttttgaatatctg gtgattgtaggcaatcacaccacagttaatggcctggcaaataatgataagcgtgtgataaacctcacaaaaatacgcctagaagtg gtgatatcgctagttttgtgtcctttcgatttttttcgtgttgcttttttctgtggataaacctttgaactttacattttgttttcaga

### col-19p

acgtaccattattcgagacaactgtgagtataaatttttggaaaaatattggatagtgaagctcataatattcaacaatgtttgtggctgctt ctgcaatctcattcaactgagctttcattgaacttgaagatctgatcacaaaagcgattgcactctgagcttcatcaaaattgtcgtccgatt gttgcagacaagaaatatgctcacaatgagcaccactgtacccaacatcacaaatgcagcctgcgttgtctggagtcgtggttccgcc attgagacaaagaggaatctcgcagttaccgccggtgaaaaattcgccacaataacatccctgaaaaaatgtagtgctattagtaatc tcatacatataaagatgacattttcctacatatgtataaaaattgcaaaaagtgaaacttcaaatatccatattctaagactttttttcaatctg ctccaatttctcaaatctgtttaaatttttgacaaatgttaatcagttttcggctatttttgatgtcacttggtcttagagcttatatagcttaacagt caaaaaagttacgcacttggaacaaaaaatgctctaaaatttccagataagaattggaaaggttttgatttgttcctttcatgaattgtggtt ttcaattgtttgttttaaaactattttcaaaaatgcagcttgcgatcattcatgtgttgttcgtagtttgtgacgtcaattttaaaatcttcaagaaa tacttgcatttctataaaattgatttgaaattttgaagcctaaattactgtcaaaaattttgattcgaaggttttctaaaacaatgagtgatttcg ggcgttatatcaaatatccaaatatataaaatttcataaaactactataaactatatgctagaaactcgtatttatactagaaatactagaa actagaaactataaaaacctggaacatcaaataaatttaatgtcagactacaaccctttaaactacataccaacgatttattagtttgag cttcaagaattttatattcttaagagggacgcacaaaaaatcactcttcacactatttctttgttgaatagaggtcagcggcttttaaaaatta taatttgaaaaaaaattgggactttttcaaaaaaaaagcttccaaacgtccctattaggaaatgtaaacttattcccaaaaaattaaaaa ttccagagaaagtagacaaatttcagaaaacttaccgcgctacatctaatactttctcataggtttttttattgggaaactgatgaaaattatt tgaattcataataaaataaattcgtatttagcatttgaaaatttgcaccaatgtattatttaattttttttttcgaaaatttaacgcattttctctctaa aaactcgaaatttagtgtgttctaaacaacagtaagcatacaatacctttgttcaaaattgacgtgctttctgaaccaatatggttagtttca aaaatttttgtattataggatagaaatatttggaaataaattttaaaaccaaacttatgcctttctcttttagtatcccagctaggtaatattttag tatttgcccaaatccttgaagtaaggagtatataatttttgaaaaacaataaaactccagataattcatagttttttctcgaaagaaaattttt gagattagttattgaacttcatttttgaacattattcgttgaaaaacactcgctttgtctttattttcaaaaaaattccgatttccccaaccagaa aaaaaaacagatagaagaaatttctccttaaatttcattgtccatctctcttggaaacacattatctatcaaatgaaaaacgcattttttttcc tggcagaaaaatgaaattggttagattacactggttaggtttgaaggtgtaactttcgctttctcagcaactttcagtataaaaggaaacg gtcaccatttagaaagacatcagttcatcaac

### myo-3p

tgtgtgtgattgctttttcacaatcagtgttttcaggattatgtgatgaactagatcttcaagtttcgttacatttcatatgttttcggaactcacga agtacatattgggtattgtgctcaaaaaattcagcaatcagcttcgctccgctgactttagaacccaaaaaaatagtatggccaaactg actgtgttacgatcatttcaatttttcaatacatatttaagatttctaagagtaagaaggtcaaaaactgttctggaatacatatatatttttcag gttacaattagtcaaaaagtgcactgaaatatacgttttaatttcacgaataacccaattagttcaatgtatttttggtcaaccaacgttaaa gtttggcttccaaccaattatcatttctgatcaaccacaatgttttttctttatctgcaagttaattttttatttttatccagatgtttggcatatttttca attcttcactagcgcccacttcttgcacttccggcgccctgaatctaatgcatctgttgcaagaattgaaagaccaatcaacacattgttttc ttcacgagatactgaagaaaatgaataaaaacagagaaaaagagccatgtgattagtgacaactgttgctaacagataccatagctt ggacttggtacgtgatggcaacgtatgggtcaacaaaaatgattgcagagggggtgcaaaacagtcaagtcgagaaaatatgaaa aacagaaaacaaagaacagaaaaatgggtttgagagtcagtataatttataaaagaaaaattgtacatagaaattaaccatttttgta gaagaagttatttttcaagcatcgttaaaaattattcaaagcaccttatttcatatttaattttaaacatggttaaatgaacaacacggtgcg caatcaggaaaacttgaaatctgaaactgttgttgtgatcttcttcgcaactgttcagatagcactagtgtaatgttaagagtgcgcgaata taatggaatataatggatcacacctcctgccatcaggtaaacgtctctgttatcacatatttccaactattaaatttttaccttttacagttttac atttttttgaaaaaagtaactttttgtcttcaaaatccctgacgaaaatatcaaatattttaatcgagactgcagaggaaccgattgatgattt ggaaaatccagctttacctgtgtaagaactgaaaagtttcataaccctagggtattcccagttacattccccactggctaacaatagcac ccagtttttcatcacctttcttcaaatttctcggcgatttgttaaaaacaaaatttgtgtcccttctctgatatctctatgtctctaaacacaagttc atcggaaaacgaaggagggtaggtgttggttgggctcccgaagtgaaaatagaagagcaagaatagaatattagagagagagtg cagagagggcgggatagctcccgggattccgttttcttcttctttatcttcaacgatgatgtgtgtgcgtgttgtatagattctgttgctccccc acaactcgctccgaaggctcaatacaattcaattgatattggaggagagcctaccggagtgggaggataagaagaaacataagaa gaagaagaagaagaagcatgcttctggtttttgatgctatgaaaacggcacaaaaagatgattgaggtcccttttcaataccttctctcat ctttcaaatcccattgaaacctaaaacttctcaccacgctttaccattgttctccaaaaacttatagcaatgtctataacttttttatctctgaaa agcagtgttccatttttctttttcctattttatttcaattgtttctcacatttcgtttggattctttgcttgtcaaccagcttcttcttccacttttaccgtcta attttcagggcagggagccatcaaacccacgaccactagatccatctagaa

### rgef-1p

aagactaattttcgattaacccgtaggggtgcaagactaatagagactgcaagactattagaggctgaaatactaattttcgtatgctca ataattttggaaattggcctattttttgtagaaacttgataccgtttaaacaaggaaaaatacacactttttaatatttcatcaataatttgaac gattttgtgattttaagttcaatttgcccaaaaaaagacaatttttctgacgttaccgcaaagtaatgctggtcaggcaaaaagagcggtg caaaaatataagagactgcaatactaatagaggaaatacggtaattgaattttagtgaaacttgcgacagttttcctcatttttttgttattcc ggcatacgagtgtggcatacgagtgaggtcatcttttgtttcttccgtttcttcatcgcttttgaaaaaaaatgttgaagaaaacttcgtagcc gatagccggataatgtttggcacggcgttccaacatatatcattggaaattttttaagatttttgcgaaaaatcacattcttcacgatgagaa cacgttattgaaggaatataaatcaagaataacatatagttatatttctctattactttaacgttaaatatgagcaaatttgagcattttgattg cgatgaaaagcagaatcgagtcaactgaaatccgttcaaaaacctaagctcggttcgcccgaaagtcgatttttccaatagccgaac agcactgcttctttattttccaggtatttttggggtgtaagaattatatcttctttgtttgggtatttacgtttccgaattccctttcttgatctgtcgcta tttttcagaaaccagaacagtttatcctttttcatgagaacctgaatagctcaaaaacccgtttaatttcttttcatcatctcatttgaaactttcc tagtcttcttaatgatttccaggctcctcctacatttgtatctcaagatgcatacactttatttccgctccaattcgaatcatctacttctcttttttctt caactttctacttttttccattcttttctttcaaatgttcgacgtcttcaagtgactgctcttcattctcctcatttttctcgaatttcattcttgtctctttttt gctcaaaaatggaaaatgaaagtgacgtgagatttggacggcgggacacgggggcagtagaagcagcaaaaaggagagaaag aggacacaaataagaagaacgaattcaaaaataagcggagaggagctatttccgtcaattctacctccccaatcttcatcaattcggc tcaattgaatgacgtcacagggagataaaccgtttggatgagcgccgtccatcactgacgccatcccgtttgggacaagaaaagaga aaaaagagcacaaagtttttggtgacggatcttgtcaatcatatgaaagttgttctgattgattgtcagtttttttcctacttttttggattctatcc acttctgaacttttgacaagtttcaaactttctgaatcattttctatgcattttcctggaattctttttatgtaaaatatgaaatagaatgtttttgaat tcaagttctgcttttttcttcttttttgttctgttcgggcttgggtatgctttttttaaaaattattttgcacatcgaccaataagtgcgcaacttataaa attaatttatttttgttaatttttgaaatacttgtattgctttaagtgatctgacctcgccctgagctttccacgtagttatcaaatacaaatcctcc aagggtaacgtacctatattactgatctttataataactttatcacctgtccagttccagaggaattctgttaagcttataggcacagaagg agtcatttctgctggtttttaatgatccaaatctttatttcaagtaaaaaactgaacacttgcgaataaaactatcagattaaccattcacca aaaatgtgtttgaatctaaaactttctcagtattccaaatatagaaataaataaccacgacattgctaaaatctgtctgaattgtgtactcctt accgtgaagtaataatggatataatgaatcgtttgaaatgaatgatcagcacatttttggtgaaagatcacaaataaggaataagcgac ggaaaataaaacgattatttcggatcaaaaatttgttgaagatcatatacactcgagaccaagattattctagacaattttcaaattggctt ctttgtttgcaaatcttcataataatctgtgaagtttggaaatttgaatttttaatcttttttcatagattatagtttttatttctttgcaaaactatattaa aaaccgatgcattgttttagggaaattaatgagccttttgttcaacactaaaaacaataaaattaaaattttggcttcatcatttgacctttttt aagttcgaaaacttttctcgtatttctgaaccgccaattttttcacacatctctagacttttggtgcccgttccagaaagttaagtaattgctatt ctaagaaagttctcaacattgttttttagttctgattgaattctgatgttccaggaatatattttaaatttaatattctttgcactatttctatactaac taaataataaatagtctaagtatgttcaatgagaacaaaaaagctctatctatatttttctatcctatttcatttatatccttttcattttgaactca ccagcttctcttcttcttcttgtccatatacttcttaaaagtctcgaacttctctctctcactctccatcaatttctttatggctaccgcttgcgctctg ccgcttccgaagaaagaaggcgttggcagagctctcaattctattttttctcgccgtaggctatcgatttctcagcttctcttctctcttcctca ctccaccaccaccacttcgggcttcttcttttcttcattttcgtcttcttcttcatcctcttctttttttttcagttccccctcactcctccccttatatgcg cgtgcgagaggggtgcaaaagcagcgcgcatccgagaattgaaacgagaaaacggagacgcagcagcagttcgtcctcagaa aaatagccagtaaaaagagaaaaagatagagagagacctatcgcattttattttcaaattgcaattcctgcaattcatgtgtgcgtgtgc ccattttcaattcttcccgttatttttcagaccattaccaacgttctttctagatcttgagacaatcttccttctgctcaatcgttcgtcgtgaagac ggcatcgacgac

### vha-6p

gcatgtacctttataggtgcgctctacgcaattcaccagctgaacaatggagttgagcctaatgtaactaaaaatttatttgaatgctttac aaaaatattatttcagatcttcgagatcatgaaaactatcaaacagcagcgccctggagcaatcgagtcgttcccacaatattcaggtgt atatgcaatcgttttagactacatttcggtaagttgctacttcagagataaactgtaattattttaaatttcagcgcaaacgcggaggaaag tctgatcctgttaacaaatacataaatcgtttcctcgctgatctcacggagatattgccagcatgctcaacgttgccagtgatgaatccaa gcacagaactgcattaagtatactatttattactcgatacttttgttcacataggttttttaaatcatattttatgcatcatttatcatattcaatgca tcattcatatcatagtcaataaaaaggttgatttctcatgttctggtttcaaatgctgactttggtaaaaagaacgcgtgcctgcctattgcct atcttggcattttctcgataaattttaaaatgtaggttcgatcttatgagatttgtagtcaaaagagctcatatgtattcaggtaggtctggtag cgagaccaacttaatagcatgacaagcattttcaatttgccctggagcgcaattggttttttattcgaaaatcgcacatttctgtttccccata atataaaatttccaggacgatatatattacattcttcacaaaatattgcattacagacaccgacaaagaatctccacctgatatgaaaac aatgagccaacaatgttatctgtattgccaccacccacatttcctagtcattcagtatatattgtttcaattgaatcattgcaggtatatatcga attgaacttgtaaggcttcatcttcatttctcaatacatcatccatcattccagagcagctccggccacacaaaaattggtggcggtctgat attgataatcgacttctttgacgtgcctgacggagcagcaaagcggagcactgataagacaatgaagaactaaaaaattgtcttcggtt ttcagtctttagttctgcagcactttattttttgtttctcctatttttccgcattttcctaactttctgatgtccatttcaaatgatttttgttataaaattgttt aatttcagggcgactaaaacctaccaaaacccataaaaa

### dsup

RNAi GCTTCAACCCATCAGTCTTCTACGGAACCATCATCGACGGGCAAGTCAGAAGAGACAAAGA AGGACGCAAGCCAGGGATCTGGACAGGATAGTAAGAATGTGACAGTCACAAAGGGAACAG GCTCTTCTGCAACTTCTGCGGCGATCGTTAAAACTGGTGGGAGTCAGGGAAAGGACAGTA GCACAACGGCTGGATCGTCGTCTACGCAAGGTCAAAAGTTTTCGACAACTCCGACTGATCC TAAGACATTTTCAAGTGATCAGAAAGAAAAGAGTAAGTCACCGGCAAAAGAGGTTCCATCG GGCGGAGACAGCAAATCTCAAGGGGATACTAAAAGCCAGTCAGATGCGAAGTCATCCGGC CAGTCCCAAGGTCAATCAAAGGATTCTGGAAAGAGCTCATCTGATTCATCAAAATCACATA GTGTGATAGGTGCCGTGAAGGATGTTGTGGCTGGGGCTAAGGACGTAGCCGGGAAGGCT GTAGAAGACGCCCCGTCTATTATGCACACCGCCGTTGACGCCGTTAAGAATGCTGCTACTA CAGTTAAAGACGTTGCCTCAAGCGCCGCTTCCACAGTAGCCGAAAAGGTGGTGGACGCCT ATCATAGCGTCGTAGGTGATAAAACTGATGATAAGAAGGAAGGAGAACATTCAGGAGATAA GAAAGATGATTCTAAGGCAGGTTCAGGATCCGGTCAGGGTGGAGATAATAAGAAATCCGA GGGTGAAACGTCAGGACAGGCTGAGAGTTCATCGGGAAATGAGGGTGCAGCACCTGCTAA GGGACGAGGCAGAGGGAGACCACCTGCTGCGGCCAAGGGCGTGGCAAAAGGAGCAGCC AAAGGAGCGGCTGCATCAAAGGGTGCAAAATCAGGCGCAGAGTCGTCTAAAGGCGGCGA GCAATCCTCGGGTGACATTGAAATGGCTGACGCAAGTTCTAAAGGTGGATCAGATCAAAGA GACTCAGCTGCTACTGTGGGTGAGGGTGGAGCTTCGG

### Lifespan and thermotolerance measurements

Lifespan assays were performed on standard RNAi plates maintained at 20 °C, following previously described procedures (Castro Torres et al., 2022). Beginning on the first day of adulthood, worms were transferred onto plates containing FUDR (100 µL of 10 mg/mL FUDR). Animals were scored every two days and monitored until each worm was either recorded as dead or censored. A worm was considered dead if it failed to move after gentle stimulation of both the head and tail using a wire pick. Individuals were censored if they exhibited bagging, extrusion of intestinal tissue through the vulva, desiccation along the edge of the plate, or other causes of death not related to aging, and these animals were excluded from further analysis.

For thermotolerance experiments, worms at day 1 of adulthood were moved onto pre-warmed NGM plates with EV bacteria, moved into a 34 °C incubator, and assessed for survival at two-hour intervals. Worms were recorded as dead if no movement was observed after stimulation with a wire at both the head and tail. Animals that showed intestinal leakage, bagging, desiccation along the edge of the plate, or other non-aging–related causes of death were classified as censored. Statistical analyses for all survival experiments were carried out using the Log-Rank test in GraphPad Prism 10 software.

### Thrashing assay

For all motility assays, thrashing behavior was measured on days 1, 5, and 9 of adulthood for most strains, with the exception of the *sur-5p::dsup* strain, which was assessed on days 1, 4, and 7 to avoid survivorship bias in these short-lived animals. Animals were grown on EV plates beginning at the L1 stage. Worms were collected using M9 buffer and transferred to an NGM plate without bacterial food. Ten-second video recordings were then captured using a Leica M205 FCA stereomicroscope. Thrashing was scored manually, where a single bend of the body to one side was counted as one thrash. The total number of thrashes occurring within the 10-second recording period was then determined.

### Mitochondrial imaging

High-resolution mitochondrial imaging was carried out using either a Leica THUNDER widefield microscope (muscle and hypodermal mitochondria) or a Leica Stellaris 5 confocal microscope (intestinal mitochondria) as previously described (Kim et al., 2025). The THUNDER system was equipped with a 63×/1.4 Plan ApoChromat objective, standard GFP filter set, a Leica DFC9000 GT camera, and a Leica LED5 light source, and was operated using LAS X software. The Stellaris 5 confocal microscope was configured with a white-light laser source, spectral filters, HyD detectors, and a 63×/1.4 Plan ApoChromat objective, and was controlled through LAS X software. For imaging, animals were transferred into M9 buffer on a glass slide and imaged within five minutes to minimize mitochondrial fragmentation. HistoBond microscope slides, which have a positively charged surface, were used to improve immobilization and reduce worm movement during image acquisition. Each experiment included three biological replicates, and a representative image was selected for presentation in the figures.

### Oxidative stress assay

Oxidative stress assays were performed using RNAi plates containing 6 mM tert-butyl hydroxide. At the L4 larval stage, worms were transferred onto tert-butyl hydroxide-containing plates with EV bacteria and FUDR and maintained at 20 °C. Animals were scored for survival approximately every 6–12 hours until all worms had died or been censored. A worm was considered dead if it failed to move following gentle stimulation of both the head and tail using a wire pick. Animals were censored if they exhibited bagging, extrusion of intestinal tissue through the vulva, desiccation along the edge of the plate, or other causes of death unrelated to oxidative stress. Statistical analysis of survival was performed using the Log-Rank test in GraphPad Prism 10 software.

### DNA damage survival assay

DNA damage tolerance assays were performed using UVC irradiation in synchronized *C. elegans* populations. On day 1 of adulthood, animals were transferred onto NGM plates containing EV bacteria with approximately 20 worms per plate across six replicate plates. Worms were placed outside of the bacterial lawn to prevent bacteria from shielding them from UV exposure. FUDR treatment was not required in this assay as embryos produced after UVC treatment are not viable.

For UVC exposure, plates were irradiated with 1 J (1,000,000 μJ) of UVC using a UV crosslinker. To minimize variation in radiation exposure, no more than three plates were placed in the crosslinker at a time and were arranged in a single line, avoiding placement near the walls of the chamber. After irradiation, worms were kept at 20 °C and scored after 24 hours. Animals exposed to UVC often exhibited reduced motility; therefore, animals were considered dead only if there was no detectable head movement or pharyngeal pumping. In cases where worms adopted a curled or rolling posture that obscured the head, animals were gently repositioned to confirm whether movement was present in the head.

After the initial scoring, worms were subjected to an additional UVC treatment of 0.5 J (500,000 μJ) and returned to incubation at 20 °C. During incubation periods, worms were placed back onto the bacterial lawn to reduce starvation, as UVC exposure can impair their ability to move toward food. Survival scoring and subsequent irradiation steps were repeated daily beginning on day 3 of adulthood until all animals had died, typically over a period of five to six days.

### Seahorse analysis

A synchronized population of worms was removed from plates using M9 buffer and transferred onto an NGM plate without bacterial food. From this plate, approximately 10-15 adult worms were carefully pipetted into individual wells of a Seahorse XF96 microplate, ensuring that no progeny were transferred along with the adults. Basal oxygen consumption rates were measured using a Seahorse XFe96 Analyzer with an XFe96 sensor cartridge. The assay cycles consisted of 2 minutes of mixing, followed by a 30-second waiting period and 2 minutes of measurement. Non-mitochondrial respiration was determined by the addition of 50 mM sodium azide, following previously described methods (Haroon and Vermulst, 2019). Oxygen consumption values were normalized based on the number of worms present in each well.

### Brood assay

For brood size assays, ten animals at the L4 larval stage were placed individually onto separate plates. Every 12 hours, adult worms were transferred to new plates, while the plates containing laid eggs were kept at 15 °C for 2-3 days to allow progeny to develop. After incubation, the number of live offspring on each plate was counted, and these values were summed to calculate the total brood size for each worm. Results are presented as a SuperPlot (Lord et al., 2020) generated in GraphPad Prism 10 software, where each individual worm is shown as a single data point, along with lines indicating the median. Statistical comparisons between groups were performed using nonparametric Mann-Whitney testing.

### Desiccation assay

For desiccation assays, adult worm expressing *vha-6p::polyQ44* were transferred onto 500 mM NaCl salt plates on day 1 of adulthood. For each condition, at least 10 worms were placed onto NGM plates lacking bacteria that contained 100 mM sodium azide (prepared in M9) to temporarily paralyze the animals. The worms were arranged in a line and imaged using a Leica M205 FCA automated fluorescent stereomicroscope controlled by LAS X software. The microscope was fitted with a standard GFP filter set, a Leica LED3 light source, and a Leica K5 camera. Three biological replicates were performed for each condition, and figures display one representative image from these replicates. Images were captured at 0, 1, 2, and 3 hours after exposure to the salt plates. To quantify polyQ foci, the number of visible foci per worm were manually counted by eye. Statistical comparisons between groups were performed using Mann-Whitney testing in GraphPad Prism 10 software.

### RNA-seq

Animals were synchronized via bleaching and L1 arresting as described, then grown on EV RNAi plates. ∼200-300 L4 animals were then manually manipulated onto fresh EV plates. For CAHS, MAHS, RvLEAM, and DSUP expressing animals, array positive animals were manually picked using the red fluorescent marker. For wild-type N2 controls, ∼200-300 animals were also manually manipulated onto new plates to ensure that manual manipulation did not contribute to transcriptional differences. Animals were then placed in a 20 °C incubator to recover for 1 hour prior to harvesting. Worms were harvested using M9, transferred into 1 mL of TRIzol Reagent, and subjected to three rounds of freeze/thaw cycles using liquid nitrogen and a 37 °C bead bath with 30 s vortexing between each cycle. After the final thaw, chloroform was added at a 1:5 ratio of chloroform:TRIzol, and RNA was separated by centrifugation in a heavy gel phase-lock tube (VWR, 10847-802). The aqueous phase was combined 1:1 with isopropanol, and purification was performed with a QuantaBio Extracta Plus RNA kit (95214) per the manufacturer’s instructions.

Library construction and sequencing were performed at Novogene using a paired-end workflow, poly(A) selection, first-strand synthesis, and Illumina NovaSeq6000. Three biological replicates were analyzed for each condition. Pre-processing to trim low-quality reads and adaptor sequences was performed with Trim Galore v0.6.7-1 (Krueger, 2025). Reads were aligned to the WBcel235 genome using STAR-2.7.3a (Dobin et al., 2013), and a raw count matrix was generated using Subread v2.0.3 (Liao et al., 2019). Differential expression was calculated with DESeq2 (Love et al., 2014) and sva v3.54.0 was employed to correct for batch effects and non-biological sources of variation (Leek et al., 2012). Gene ontology enrichment analysis was performed using WormEnrichr (Chen et al., 2013; Kuleshov et al., 2016).

## Author Contributions

A.E. designed and performed all *C. elegans* experiments and prepared the manuscript and figures. A.A., D.H., E.A.R., A.J.C., and A.J.H. assisted with lifespans and stress assays. J.K. performed DNA damage assays and assisted with mitochondrial imaging. M.A., S.P., R.A.B., T.J.L., V.K., M.P., V.S., B.V.T., and J.D.K. assisted with stress assays. A.A. assisted with all strain construction. N.D. assisted with mitochondrial imaging. P.J.M. assisted with experimental design and supervision. G.G. and R.H.S. designed and assisted with all *C. elegans* experiments, preparation of the manuscript and figures, and performed all supervision for work. All authors edited and approved the manuscript.

## Data Availability

All data required to evaluate the conclusions in this manuscript are available within the manuscript and Supplementary Materials. All strains synthesized in this manuscript are derivatives of N2 or other strains from CGC and are either made available on CGC or available upon request. All raw RNA-seq datasets are available at the National Center for Biotechnology Information Sequence Read Archive repository accession: PRJNA1550449. All code used to analyze datasets included in this work is available on the Sanabria lab GitHub repository: https://github.com/SanabriaLab/Entin-et-al.-2026.

## Acknowledgements

A.A., A.J.C., A.J.H., and M.A. are supported by the T32AG052374 from the NIA. J.K. is supported by the USC Provost Fellowship and Leonard Davis School of Gerontology Biology of Aging PhD Fellowship. A.J.C. is supported by the Diana Jacobs Kalman/AFAR Graduate Student Awards. A.J.H. is supported by the National Science Foundation GRFP DGE-1842487. E.D. and T.W. are supported by the COMPASS CIRM grant. P.J.M. is supported by the Impetus Grants 014746-00001 and the Baxter award. G.G. is supported by T32AG052374 and R01AG079806-02S1 from the NIA. R.H.S. is supported by R01AG079806 from the NIA. Real-time oxygen consumption rates (OCR) of *C. elegans* were measured by Dr. Hemal Mehta using the XF96 Analyzer (Seahorse Bioscience) at the USC Leonard Davis School of Gerontology Seahorse Core. Some strains were provided by the CGC, which is funded by the NIH Office of Research Infrastructure Programs (P40 OD010440). Some gene analysis was performed using Wormbase, which is funded on a U41 grant HG002223. Some studies were conducted at the Marine Biological Laboratory through the Biology of Aging course.

---

**Fig. S1.**
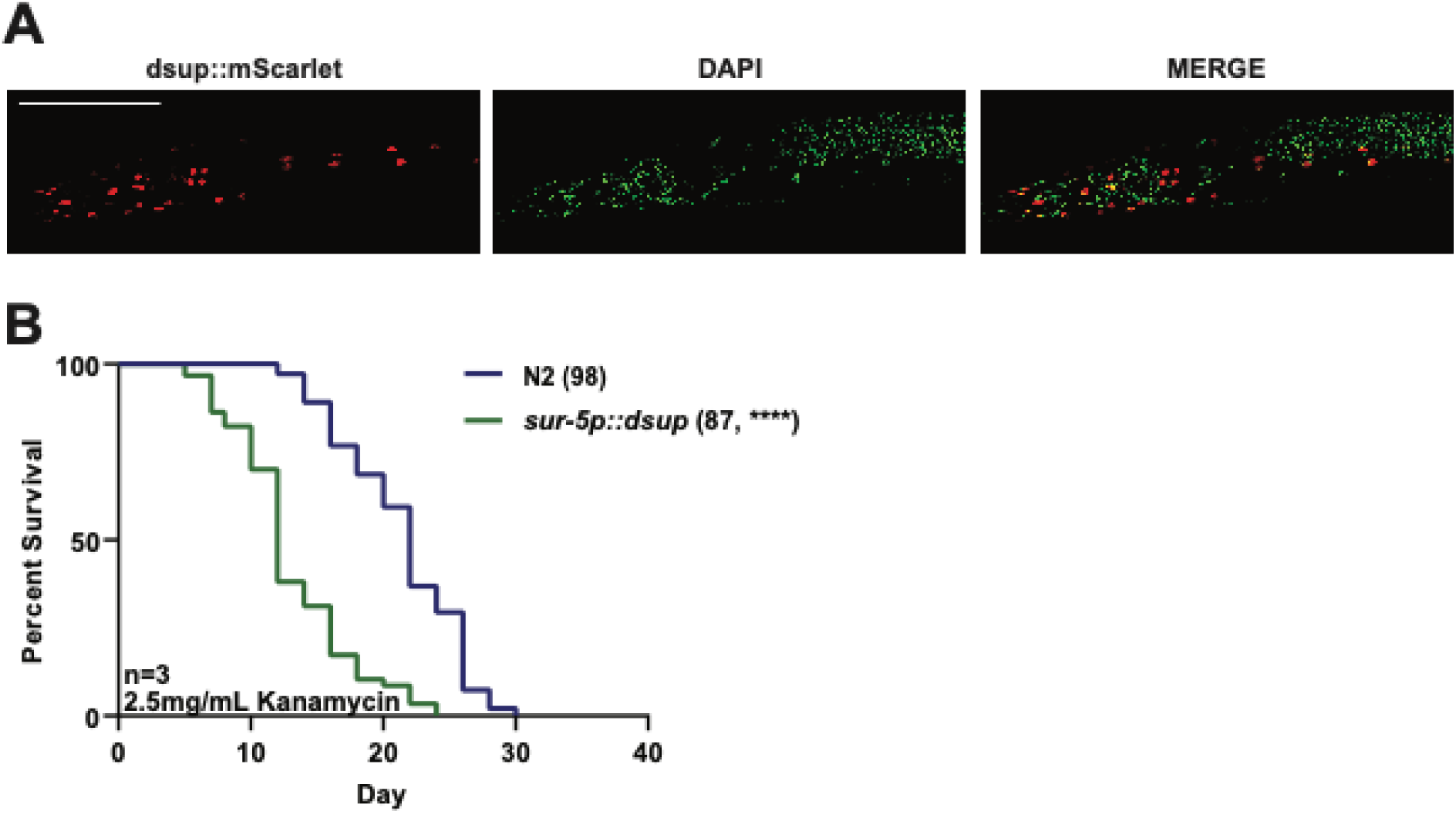
Somatic DSUP localizes to nuclei and reduces lifespan independently of bacterial pathogenesis. **(A)** Representative fluorescent images of animals expressing DSUP in somatic tissues (*sur-5p::dsup*). DSUP fluorescence is shown in red, DAPI nuclear staining in green, and the merged image demonstrates nuclear localization of DSUP. Scale bar is 100 µm. **(B)** Lifespan of wild-type N2 and *sur-5p::dsup* animals maintained on kanamycin-killed bacteria. Numbers in parentheses indicate total animals scored. Statistical significance was determined by Log-Rank testing. Statistical information for all replicates is available in **Table S2**. Note: the lifespan data was collected and analyzed together with data from **Fig. S2D**. **** = p < 0.0001.

**Fig. S2.**
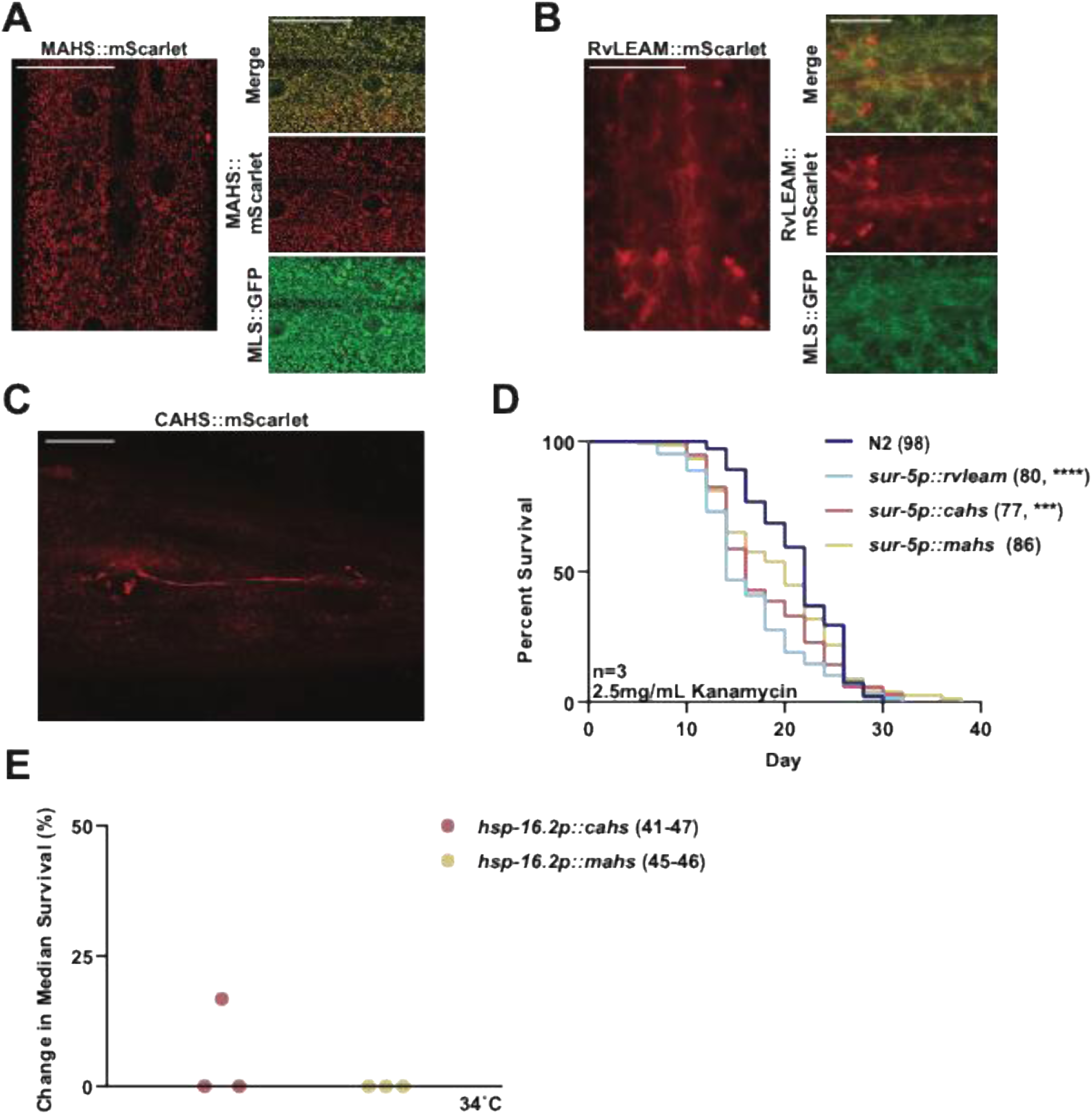
Subcellular localization of RvLEAM, CAHS, and MAHS in *C. elegans*. **(A)** Representative fluorescent images of animals expressing MAHS::mScarlet shown in red, mitochondrial matrix-localized GFP (MLS::GFP) in green, and the merged image demonstrates colocalization of MAHS with mitochondria. Scale bar is 20 µm. **(B)** Representative fluorescent images of animals expressing RvLEAM::mScarlet shown in red, MLS::GFP in green, and the merged image demonstrates predominant mitochondrial localization of RvLEAM. Scale bar is 10 µm. **(C)** Representative fluorescent image of animals expressing CAHS::mScarlet, demonstrating diffuse cytoplasmic localization. Scale bar is 20 µm. **(D)** Lifespan of wild-type N2 animals and transgenic animals constitutively expressing RvLEAM (*sur-5p::rvleam*), CAHS (*sur-5p::cahs*), or MAHS (*sur-5p::mahs*) in somatic tissues. Animals are grown on bacteria treated with 2.5 mg/mL kanamycin. Numbers in parentheses indicate total animals scored. Statistical significance was determined by Log-Rank testing. Statistical information for all replicates is available in **Table S2**. Note: the lifespan data was collected and analyzed together with data from **Fig. S1B. (E)** Change in median survival during thermotolerance assays of animals expressing CAHS or MAHS under the heat-shock-inducible *hsp-16.2* promoter. Animals were exposed to 3°C, and each point represents one independent biological replicate. N=34-47. *** = p < 0.001; **** = p < 0.0001; ns = not significant, p > 0.05.

**Fig. S3.**
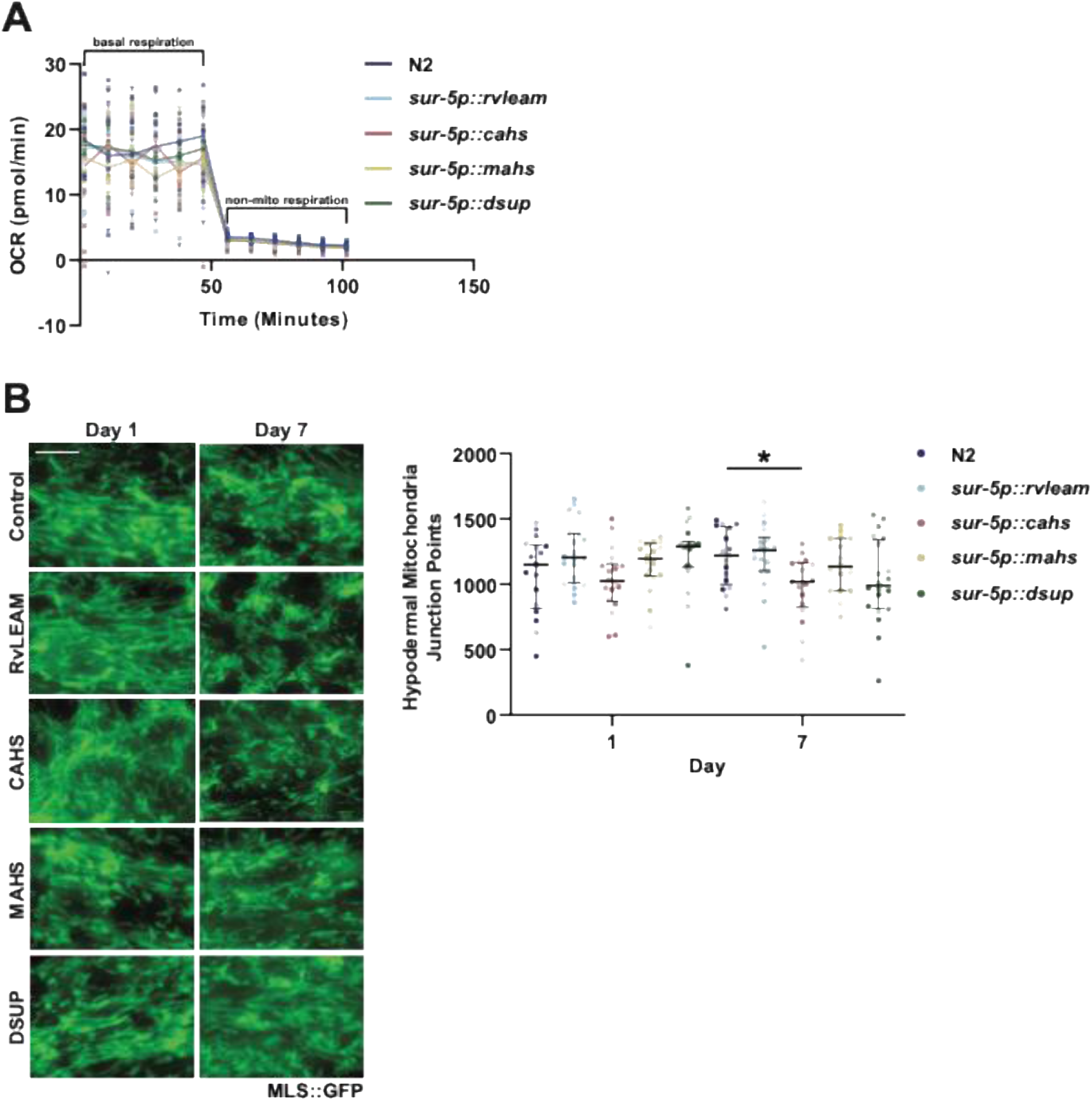
Somatic RvLEAM, CAHS, MAHS or DSUP expression does not significantly alter mitochondrial respiration, and only CAHS expression affects mitochondrial morphology in the hypodermis. **(A)** Basal oxygen consumption rate (OCR) of wild-type N2, *sur-5p::rvleam*, *sur-5p::cahs*, *sur-5p::mahs*, and *sur-5p::dsup* animals. OCR was measured using a Seahorse XFe96 Analyzer and normalized to the number of worms per well. Sodium azide was added to determine non-mitochondrial respiration. Data is presented as a SuperPlot with each individual dot representing one animal, different shades indicating each independent biological replicate, and line indicating the median at all time points. **(B)** Right: Representative fluorescent images of hypodermal mitochondrial morphology (*col-19p::MLS::GFP*) at day 1 and day 7 in wild-type N2, *sur-5p::rvleam* (RvLEAM), *sur-5p::cahs* (CAHS), *sur-5p::mahs* (MAHS), and *sur-5p::dsup* (DSUP) animals. Scale bar is 5µm. Left: Quantification of images using mitoMAPR. Mitochondrial morphology was quantified by the number of mitochondrial junction points. Data is presented as a SuperPlot with each individual dot representing one animal, different shades indicating each independent biological replicate, and lines indicating the median and interquartile range. * = p < 0.05 using non-parametric Mann-Whitney testing. All raw mitoMAPR analysis is available in **Table S3**.

**Fig. S4.**
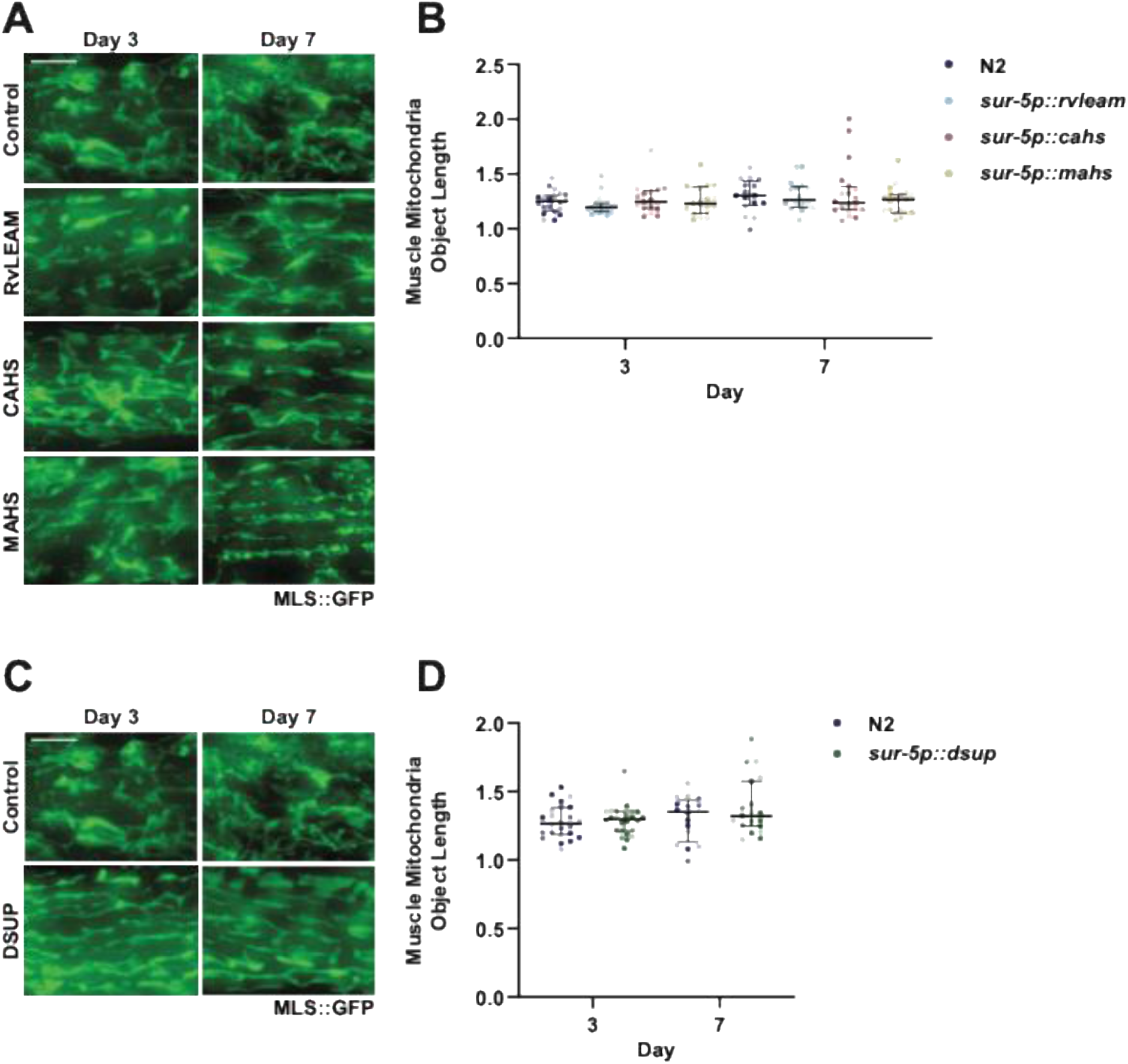
Somatic RvLEAM, CAHS, MAHS or DSUP expression does not significantly alter mitochondrial morphology in the muscle. **(A)** Representative fluorescent images of muscle mitochondrial morphology (*myo-3p::MLS::GFP*) at day 1 and day 7 in wild-type N2, *sur-5p::rvleam* (RvLEAM), *sur-5p::cahs* (CAHS), and *sur-5p::mahs* (MAHS). Scale bar is 5µm. **(B)** Quantification of images using mitoMAPR. Mitochondrial morphology was quantified by mitochondrial object length. Data is presented as a SuperPlot with each individual dot representing one animal, different shades indicating each independent biological replicate, and lines indicating the median and interquartile range. Note: two of three replicates were collected and analyzed together with data from **Fig. S4D**. **(C)** Representative fluorescent images of muscle mitochondrial morphology (*myo-3p::MLS::GFP*) at day 1 and day 7 in wild-type N2 and *sur-5p::dsup* (DSUP) animals. Scale bar is 5µm. **(D)** Quantification of images using mitoMAPR. Mitochondrial morphology was quantified by mitochondrial object length. Data is presented as a SuperPlot with each individual dot representing one animal, different shades indicating each independent biological replicate, and lines indicating the median and interquartile range. All statistical testing were not significant using non-parametric Mann-Whitney testing. All raw mitoMAPR analysis is available in **Table S3**. Note: two of three replicates were collected and analyzed together with data from **Fig. S4B**.

**Fig. S5.**
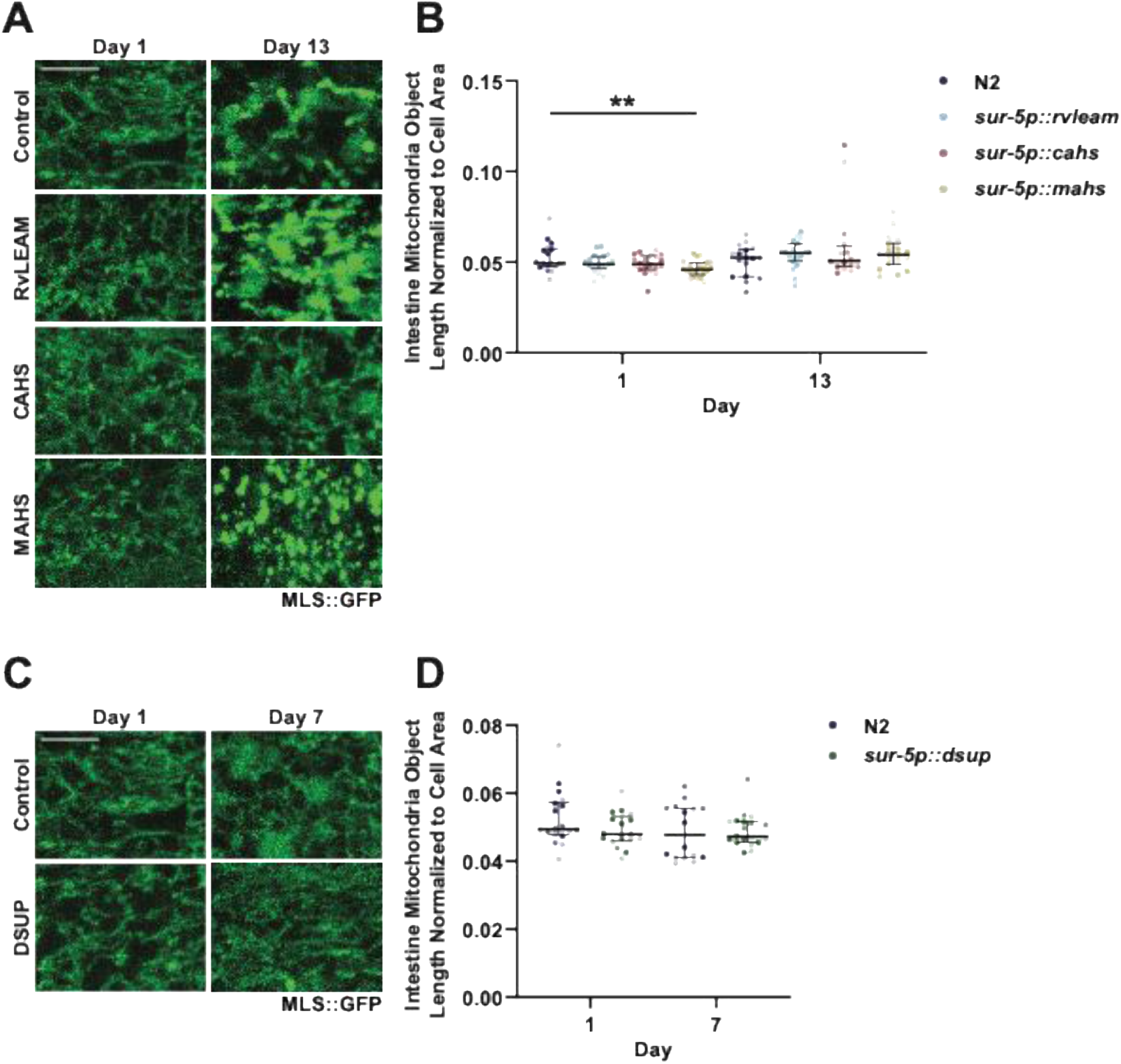
Somatic RvLEAM, CAHS, or DSUP expression does not significantly alter mitochondrial morphology, while MAHS has subtle, but statistically significant changes in the intestine. **(A)** Representative fluorescent images of intestinal mitochondrial morphology (*vha-6p::MLS::GFP*) at day 1 and day 13 in wild-type N2, *sur-5p::rvleam* (RvLEAM), *sur-5p::cahs* (CAHS), and *sur-5p::mahs* (MAHS) animals. Scale bar is 5µm. **(B)** Quantification of (A) using mitoMAPR. Mitochondrial morphology was quantified by mitochondrial object length normalized to the cell area. Data is presented as a SuperPlot with each individual dot representing one animal, different shades indicating each independent biological replicate, and lines indicating the median and interquartile range. Note: the day 1 data was collected and analyzed together with data from **Fig. S5D**. **(C)** Representative fluorescent images of intestinal mitochondrial morphology (*vha-6p::MLS::GFP*) at day 1 and day 7 in wild-type N2 and *sur-5p::dsup* (DSUP) animals. Scale bar is 5µm. **(D)** Quantification of (C) using mitoMAPR. Mitochondrial morphology was quantified by mitochondrial object length normalized to the cell area. Data is presented as a SuperPlot with each individual dot representing one animal, different shades indicating each independent biological replicate, and lines indicating the median and interquartile range. All statistical testing were not significant using non-parametric Mann-Whitney testing. All raw mitoMAPR analysis is available in **Table S3**. Note: the day 1 data was collected and analyzed together with data from **Fig. S5B**.

**Fig. S6.**
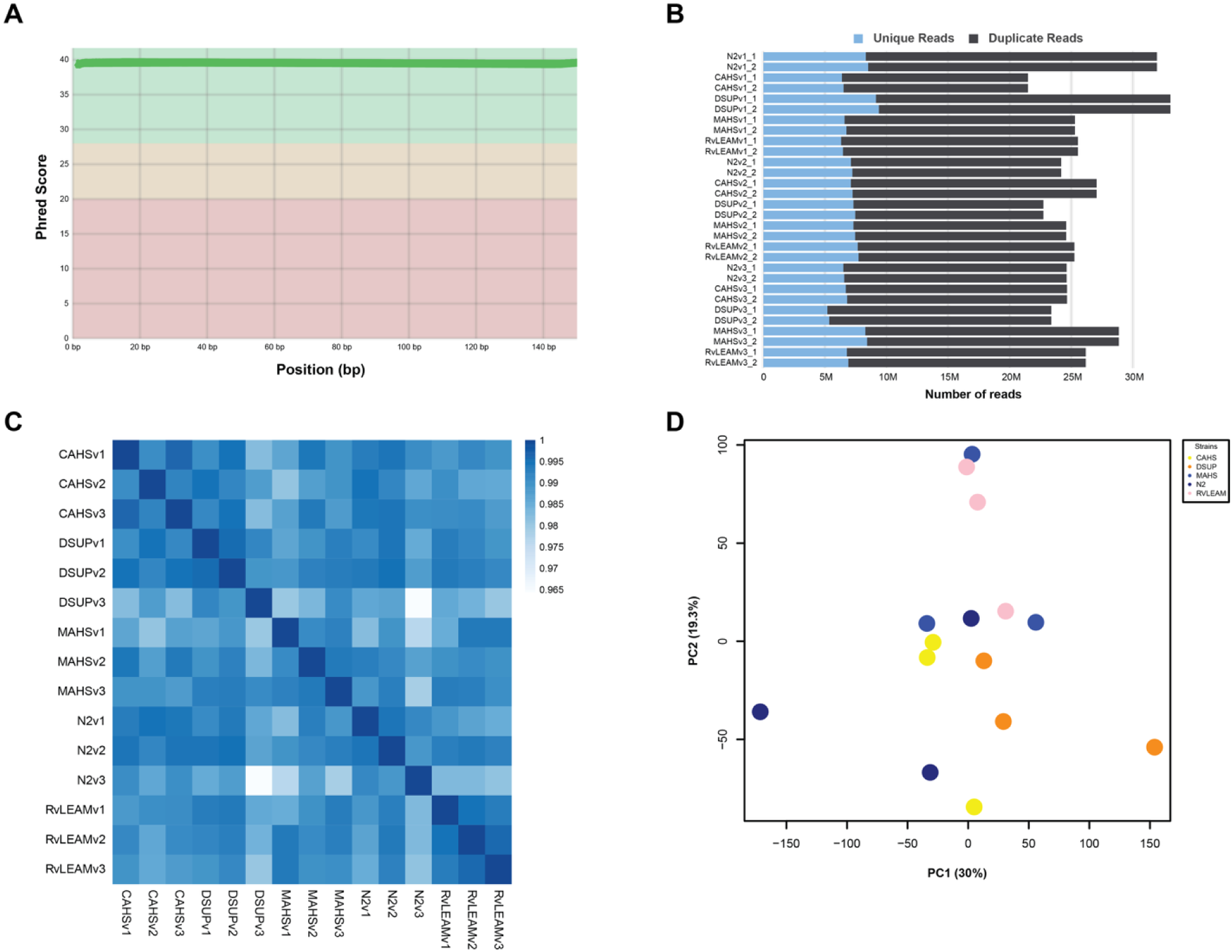
RNA-seq quality control and sample assessment for TDP-expressing C. elegans strains. **(A)** Per-base sequence quality scores across reads, showing consistently high Phred scores across all nucleotide positions. **(B)** Number of unique and duplicate reads for each biological replicate of N2, CAHS, DSUP, MAHS, and RvLEAM samples. **(C)** Correlation heatmap showing pairwise similarity among RNA-seq samples, with biological replicates generally exhibiting high correlation. **(D)** Principal component analysis (PCA) of RNA-seq samples, with points colored by strain. PC1 and PC2 explain 30% and 19.3% of the total variance, respectively, and illustrate variation both within and between strain groups.

**Fig. S7.**
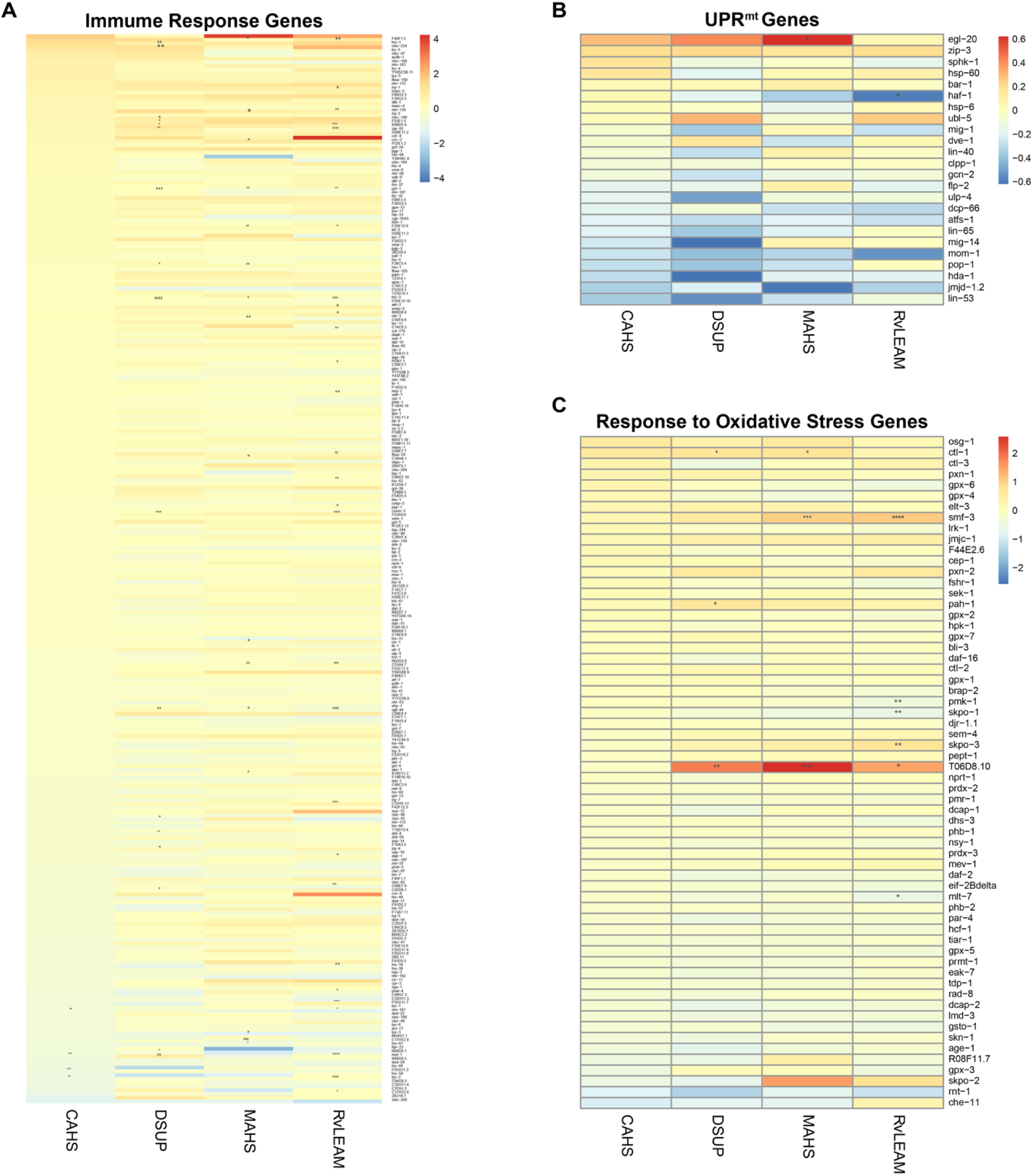
Tardigrade-derived proteins alter expression of stress-response and immune-related genes. **(A)** Heatmap showing differential expression of immune response genes (GO:0006955) in animals expressing CAHS, DSUP, MAHS, or RvLEAM relative to control animals. **(B)** Heatmap showing differential expression of genes associated with the mitochondrial unfolded protein response (GO:0034514) across the four TDP-expressing strains. **(C)** Heatmap showing differential expression of genes associated with the response to oxidative stress (GO:0006979). Colors indicate the magnitude and direction of gene expression changes, with red representing increased expression and blue representing decreased expression relative to control. Asterisks indicate statistically significant differential expression. All differentially expressed genes for RNA-seq experiments are available in **Table S4** and genes used to construct heat-maps are available in **Table S5**.

